# Translating Sex- and Disease-Informed Design Criteria from Human Lung into Engineered Biomaterials

**DOI:** 10.64898/2026.10.27.762197

**Authors:** Haley Noelle, Mikala C. Mueller, Rachel Blomberg, Nicole D. Sempertegui, Dema Essmaeil, Brisa Peña, Maxwell McCabe, Chelsea M. Magin

**Author notes:** **Corresponding Author**: Chelsea M. Magin, PhD, 2115 N Scranton Street, Suite 3010, Aurora, CO 80045.

## Abstract

Idiopathic pulmonary fibrosis (IPF) is a progressive lung disease characterized by pathological extracellular matrix (ECM) remodeling, and a pronounced sex disparity, with approximately 70% of cases occurring in males. Despite these differences, biomaterials used to model fibrosis are commonly designed to replicate disease state or stiffness alone and rarely account for biological sex. Here, we derived sex- and disease-informed biomaterial design criteria from female and male control and IPF human lungs and translated these criteria into engineered hybrid-hydrogels containing human lung decellularized ECM (dECM) and poly(ethylene glycol) alpha methacrylate (PEG⍰MA). Matrisome-focused reanalysis of our previously published region-resolved human lung proteomic dataset revealed region- and sex-associated differences in fibrillar collagens, matricellular proteins, and structural proteoglycans. Complementary characterization of human lung tissue further identified sex- and disease-associated differences in fibrillar collagen architecture, proteoglycan and glycosaminoglycan (GAG) remodeling, and tissue mechanics. Male IPF lungs exhibited the highest levels of disease-associated stiffening and fibrillar collagen remodeling, while female IPF tissue showed comparatively modest mechanical changes despite biochemical remodeling. These measurements established tissue-informed design criteria spanning matrix composition, architecture, and mechanics. We next evaluated whether tissue-derived properties were reproduced or retained following decellularization, processing, and incorporation of donor-derived dECM into hybrid-hydrogels. Engineered hybrid-hydrogels closely recapitulated sex- and disease-associated mechanical relationships, while COL1A1-associated abundance patterns were more consistently preserved than fibrillar collagen organization. Matricellular proteins and GAG-rich matrix remained detectable, although condition-dependent biochemical fidelity varied. This work establishes a human tissue-informed framework for engineering sex- and disease-specific biomaterials, highlighting the importance of defining sex-associated ECM properties and validating individual features following material fabrication to advance human models of IPF and other diseases with sex-associated differences in susceptibility and progression.

## 1. Introduction

Sex-dependent biology remains insufficiently integrated into biomaterials research, despite increasing recognition that male and female cells respond differently to biochemical, chemical, and mechanical cues.^1–7^ This limitation extends beyond cell-intrinsic behavior to the extracellular matrix (ECM), which provides critical biochemical, structural, and mechanical signals that regulate tissue homeostasis and disease progression.^8–11^ However, most ECM analyses have not been stratified by sex, limiting the tissue-level information available to design physiologically relevant biomaterials.^5, 6^ As a result, current disease models may overlook microenvironmental features that differ between females and males. This consideration is particularly important when designing three-dimensional *in vitro* models intended to capture sex-specific cellular behavior.^12^ Sex differences in ECM have been observed in healthy and diseased tissues, suggesting that females and males may not share the same baseline extracellular microenvironment before disease onset.^13, 14^ Thus, sex-informed biomaterial design requires defining both baseline differences in the native microenvironment and how those environments are differentially remodeled during disease. This is especially important for fibrotic diseases, where the ECM is not merely a downstream consequence of pathology but an active regulator of cell behavior, tissue mechanics, and disease progression.^9, 15, 16^

Idiopathic pulmonary fibrosis (IPF) is a progressive interstitial lung disease characterized by irreversible scarring, loss of respiratory function, limited treatment options, and poor clinical outcomes.^15, 17–20^ Approximately 70% of IPF cases occur in males, and males generally exhibit worse clinical outcomes than females, suggesting that sex-associated biology may contribute to disease susceptibility and progression.^21–23^ These epidemiological differences motivate direct investigation of sex-associated disease biology in human lung tissue.^24, 25^ IPF is driven by aberrant repair following repetitive epithelial injury, resulting in persistent fibroblast activation and pathological ECM remodeling.^26–31^ This remodeling is spatially heterogeneous, with fibroblastic foci localized areas of active matrix synthesis adjacent to relatively preserved alveolar regions.^26–33^ Comparing alveolar and fibroblastic-focus ECM therefore provides spatial resolution of fibrotic remodeling that is obscured by analysis of bulk fibrotic lung tissue alone.^32–34^

Although collagen accumulation is a defining feature of fibrosis, collagen abundance alone cannot fully explain the mechanical behavior of fibrotic lung tissue.^10, 16, 35^ Matrix mechanics arise from multiple ECM properties, including fibrillar collagen organization and crosslinking, proteoglycan and glycosaminoglycan (GAG) composition, matrix hydration ^36^, glycoprotein-mediated fiber interactions, and regional tissue architecture.^37–40^ Increased ECM stiffness is also a biologically active process – fibroblasts and epithelial cells sense and respond to changes in matrix mechanics, and increased stiffness can reinforce fibroblast activation, myofibroblast persistence, ECM deposition, and progressive remodeling.^9, 11, 32, 41, 42^ Matrix stiffness is therefore an emergent property of multiple ECM features that can both reflect and regulate fibrotic remodeling.^10, 16^ Accordingly, sex-associated differences in IPF mechanics may arise from broader differences in matrix composition, organization, and supramolecular architecture rather than collagen abundance alone.^43, 44^ This distinction is particularly relevant to biomaterial design because matching disease-associated stiffness without reproducing the matrix features that generate or accompany that stiffness may incompletely represent the native microenvironment.^12, 16, 45–47^ A central unanswered question is therefore whether female and male lungs begin from the same ECM baseline and whether IPF remodels those matrices through the same structural, biochemical, and mechanical programs. Addressing this question requires distinguishing baseline sex differences, disease-associated remodeling within each sex, and sex differences within the fibrotic state. This framework provides tissue-informed design criteria that capture sex- and disease-associated differences that would be obscured by pooling male and female tissues into common control and IPF groups.

Biomaterial platforms provide a unique opportunity to reconstruct defined features of the native extracellular microenvironment.^12, 48^ Human lung dECM hydrogels, for example, retain disease-associated ECM properties following decellularization and material fabrication. Hydrogels generated from control and IPF human lungs broadly preserved patterns in native tissue mechanics and viscoelasticity.^49, 50^ More recent studies further demonstrated region- and disease-specific differences in the composition of airway-, alveolar-, and vascular-enriched human lung dECM, highlight the importance of tissue source when designing lung biomaterials.^50^ Despite the marked sex disparity in human IPF, pulmonary biomaterials are typically created from pooled or sex-agnostic source material and therefore may not reproduce sex-associated features of the native lung microenvironment.^12^ If female and male matrices differ at baseline, or undergo distinct remodeling during fibrosis, an averaged material may obscure rather than reproduce the biologically relevant microenvironments.^34^ Moreover, tissue derivation alone does not establish that a dECM-based material preserves the native tissue properties: decellularization, enzymatic digestion, functionalization, and network formation can alter ECM composition, architecture, and mechanics.^49–52^ Tissue-informed biomaterial design therefore requires first defining relevant native tissue properties and then verifying that those properties remain present after material fabrication.

Our previous hybrid-hydrogel studies established a platform that combines tissue-derived dECM with independent control of matrix mechanics,^53–57^ while our recent region-resolved proteomic analysis identified regional and sex-associated variation in female and male control and IPF human lungs.^34^ Here, we integrate these complementary approaches to define sex- and disease-informed biomaterial design criteria directly from human lung tissue and translate these tissue-informed criteria into engineered biomaterials.^12^ We performed a matrisome-focused reanalysis of control alveoli, IPF alveoli, and fibroblastic foci and integrated these proteomic data with orthogonal measurements of native tissue mechanics, fibrillar collagen architecture, matricellular proteins, and GAG-rich matrix. We then translated these tissue-informed design criteria into hybrid-hydrogels composed of human lung dECM and poly(ethylene glycol) alpha-methacrylate (PEGαMA) enabling retention of sex-associated biochemical cues and independent control of matrix mechanics. Together, this work establishes a human tissue-informed framework for engineering sex- and disease-specific biomaterials and demonstrates the importance of defining the native microenvironment and validating which tissue features are preserved after biomaterial fabrication.

## 2. Materials and Methods

### 2.1. Human Lung Tissue and Study Design

Human control and IPF lung tissues were obtained under approved human-subjects protocols. All IPF specimens included in the region-resolved proteomic cohort met established diagnostic criteria according to the American Thoracic Society/European Respiratory Society/Japanese Respiratory Society/Latin American Thoracic Association (ATS/ERS/JRS/ALAT) clinical practice guidelines.^58^ Written informed consent was obtained from participants, and tissues were collected under Colorado Multiple Institutional Review Board approval (COMIRB 15-1147) or obtained as de-identified human specimens through the Lung Tissue Research Consortium (LTRC). Human lung cryosections used for tissue characterization were prepared through the Human Tissue and Cell Core, supported by the National Heart, Lung, and Blood Institute (NHLBI) grant P01HL162607. Human lung tissue used for decellularization was obtained through the LTRC at National Jewish Health.

The region-resolved human lung proteomic dataset was previously published by our group and was reanalyzed here to characterize sex-, disease-, and region-associated differences in the human lung matrisome.^33, 34^ The cohort included donors with and without the *MUC5B* promoter risk variant, with carrier frequencies ranging from 40-66% across sex- and disease-stratified groups. Additional human lung specimens were used for orthogonal human lung tissue characterization, dECM preparation, and hybrid-hydrogel fabrication. Samples were stratified by donor sex and disease status, with individual donor characteristics tracked throughout native-tissue characterization, dECM processing, hydrogel fabrication, and downstream material characterization. Donor demographics, tissue sources, and experiments performed for each donor are summarized in Table S1.

### 2.2. Laser Capture Microdissection Coupled Mass Spectrometry

Sample processing, laser capture microdissection (LCM), and mass spectrometry were performed as previously described.^33, 34^ Briefly, 5-µm-thick formalin-fixed, paraffin-embedded human lung sections were stained with hematoxylin and eosin to enable identification of anatomical regions by tissue morphology. Control alveoli, IPF alveoli, and fibroblastic foci were isolated using the Molecular Machinery Instruments (MMI) CellCut system coupled to an Olympus IX63 microscope. Multiple regions from each anatomical compartment were pooled within each donor to obtain approximately 0.005–0.010 mm³ of tissue. Following protein extraction and tryptic digestion, peptides were analyzed using a NanoElute liquid chromatography system coupled to a timsTOF SCP mass spectrometer (Bruker).

### 2.3. Matrisome-Focused Proteomic Data Analysis

Raw proteomic data were reprocessed and analyzed as previously described.^34^ Briefly, peptide and protein identifications were generated against the SwissProt *Homo sapiens* database, with identifications filtered to a 1% false discovery rate (FDR).^34^ Label-free intensities were analyzed using MetaboAnalyst 6.0 (accessed February 2026).^59, 60^ Proteins within the lowest 5% of mean intensity values were excluded, and remaining intensities were log10-transformed. Principal component analysis (PCA) and two-way analysis of variance (ANOVA) were performed using an FDR threshold of 0.10. Partial least-squares discriminant analysis (PLS-DA) was used to calculate variable-importance-in-projection (VIP) scores, with VIP > 1 indicating contribution to group separation. Differentially abundant proteins were identified using a ≥2-fold-change threshold and raw p ≤ 0.10. For the present study, subsequent analyses were restricted to proteins classified within the human matrisome according to the MatrisomeDB4 (annotations levels 1 and 2) annotations.^61, 62^ Heatmaps were generated from the indicated protein sets using Ward’s hierarchical clustering method. Protein intensities were row-scaled for heatmap visualization. Matrisome-composition analyses were performed separately using nontransformed protein intensities, ensuring that relative protein contributions were not calculated from row-scaled heatmap values.

### 2.4. Matrisome Composition Analysis

Matrisome proteins were classified into core-matrisome (collagens, ECM glycoproteins, and proteoglycans) and matrisome-associated categories using established human matrisome annotations.^61, 62^ Total matrisome abundance included both core and matrisome-associated proteins. Total matrisome abundance was calculated by summing the nontransformed intensities of annotated matrisome proteins within the analyzed sample. Relative core-matrisome composition was calculated using nontransformed intensities to determine the contributions of collagens, ECM glycoproteins, and proteoglycans across sex, disease status, and region. Each core-matrisome class was expressed as a percentage of total core-matrisome intensity, calculated as the summed intensity of the corresponding class divided by the summed intensity of all core-matrisome proteins and multiplied by 100. The contributions of individual fibrillar collagens, matricellular glycoproteins, and structural proteoglycans were expressed as percentages of the summed intensity of their respective subclasses. This approach distinguished changes in relative matrix composition from changes in total protein abundance.

### 2.5. Pathway Enrichment Analysis

Pathway enrichment analysis was performed separately for female and male datasets using Metascape (accessed February 2026). Enrichment was evaluated across KEGG Pathway, Gene Ontology Biological Processes, Reactome Gene Sets, Canonical Pathways, CORUM, WikiPathways, and PANTHER Pathway, using the full genome as the background. Enriched terms were identified using p < 0.01, a minimum count of three, and an enrichment factor >1.5. Statistical significance was evaluated using the cumulative hypergeometric distribution with Benjamini–Hochberg correction for multiple testing. Selected enriched pathways were visualized as horizontal bar plots and grouped by biological function into four categories: ECM organization and matrisome architecture, matrix turnover and remodeling-associated signaling, cell–matrix adhesion and integrin signaling, and proteoglycan and pericellular-matrix remodeling. These functional groups were assigned using established matrisome classifications and literature-supported relationships among matrix proteins, integrin-mediated adhesion, matrix remodeling, and proteoglycan-associated signaling.^61–64^ Group assignments were used to organize and interpret related enrichment terms.

### 2.6. Immunofluorescence Staining and Analysis of Human Lung Tissue

Human lung tissues were flash frozen, embedded in optimal cutting temperature (OCT) compound, cryosectioned at 10-µm thickness, and stored frozen until staining. Sections were equilibrated to room temperature for approximately 15 min, rinsed in phosphate-buffered saline (PBS) to remove residual OCT, and lightly fixed in ice-cold acetone for 15 min. Sections were outlined with a hydrophobic barrier pen and blocked in 5% bovine serum albumin (BSA) in PBS for 30 min at room temperature. Sections were incubated with primary antibodies diluted in blocking buffer (5% BSA in PBS) overnight at 4 °C. Negative-control sections were incubated in blocking buffer without primary antibody. Sections were washed three times in PBS containing 1% Tween-20 (PBST) and incubated with species-appropriate fluorescent secondary antibodies for 1 h at room temperature. For antibody panels containing goat anti-human primary antibodies, sections were incubated with donkey anti-goat secondary antibody before subsequent secondary-antibody incubation to minimize cross-reactivity. Sections were then washed, counterstained with Hoechst 33342 (ApexBio; 10 µg/mL) for 15 min, washed with PBS, and mounted using ProLong Gold Antifade Reagent (Thermo Fisher Scientific). Slides were cured overnight protected from light before imaging on an Olympus BX63 epifluorescence microscope. Primary and secondary antibodies used for COL1A1, TNC, POSTN, and associated markers are provided in Table S2.

Immunofluorescence images were analyzed using Fiji/ImageJ. Rolling-ball background subtraction with a radius of 50 pixels was applied before separating and independently thresholding fluorescence channels. Regions of interest (ROIs) were manually defined according to tissue morphology and anatomical compartment. For the lung immunofluorescence analyses presented in Figures 5 and 6, the finalized alveolar-region dataset contained 403 ROIs across 12 human lung donors, with three independent donors represented in each sex-by-disease group. These included 142 COL1A1 ROIs, 122 POSTN ROIs, and 139 TNC ROIs.

Marker-positive area was quantified within the tissue-containing portion of each ROI and expressed as a percentage of the analyzed ROI area: marker-positive area (%) = 100 × (marker-positive pixels / tissue-containing ROI pixels). The donor was the biological experimental unit, and repeated ROI measurements were accounted for in donor-level mixed-effects analyses.

### 2.7 Alcian Blue Staining and Analysis of Human Lung Tissue

Human control and IPF lung tissues were cryosectioned at 10-µm thickness and equilibrated to room temperature for approximately 10–15 min before staining. Sections were fixed in ice-cold acetone for 10 min, air-dried for approximately 10 min, and gently rehydrated in deionized water. Alcian Blue 8GX was prepared at 1% (w/v; 1 g/100 mL) in 3% acetic acid and adjusted to pH 2.5. Sections were incubated in Alcian Blue solution for 30 min at room temperature to visualize acidic GAG-rich matrix and subsequently washed three times for 1 min each in deionized water. Following staining, sections were dehydrated in 95% ethanol for 3 min, followed by two changes of 100% ethanol for 3 min each. Sections were then counterstained with Hematoxylin and Eosin. Samples were cleared using two changes of SafeClear II for 3 min each and mounted using Cytoseal 60 mounting medium.

Brightfield images were acquired under consistent illumination, exposure, and white-balance settings across experimental conditions. Images were analyzed in Fiji/ImageJ, with ROIs defined to exclude non-tissue background. Alcian Blue-positive area was quantified using a consistent color-thresholding procedure and normalized to the tissue-containing area within the corresponding ROI. The percentage of Alcian Blue-positive area was calculated as 100 × (Alcian Blue-positive area / total tissue-containing ROI area).

The finalized lung Alcian Blue analysis used 84 image/ROI measurements from 12 donors, with three donors represented per sex-by-disease condition. Repeated ROI measurements were treated as spatial subsamples, with the donor serving as the biological replicate. Results were expressed as the mean percentage of Alcian Blue-positive tissue area per donor.

### 2.7. Atomic Force Microscopy and Analysis of Human Lung Tissue

Mechanical properties of human distal lung tissue were measured using a NanoWizard 4a atomic force microscope (JPK Instruments). Lung fragments approximately (∼1 cm) were embedded in OCT, cryosectioned at 10-µm thickness, and mounted on glass slides. Sections were rinsed in PBS under gentle agitation for 5–10 min to remove OCT and tissue debris. Force spectroscopy was performed using an MLCT probe (Bruker) with a nominal spring constant of 0.01–0.02 N/m with a maximum applied force was limited to 1.5 nN and a 5-µm Z-piezo displacement. At least 50 indentations were collected per sample across five fields of view identified using 40× phase-contrast imaging. Measurement regions were selected based on tissue morphology, targeting fibrotic regions in IPF samples and morphologically normal alveolar regions in controls.

Force curves were fitted using the Hertz contact model in JPK analysis software to calculate Elastic modulus. Indentations and fields of view were treated as technical and spatial subsamples, respectively, with donor serving as the biological replicate. Elastic modulus measurements were reported as geometric means.

### 2.8. PEGαMA Synthesis for Hybrid-Hydrogel Formation

Eight-arm poly(ethylene glycol)-OH (PEG-OH; 10 kg/mol; Jen-Kem Technology) was dissolved in anhydrous tetrahydrofuran (THF; Sigma-Aldrich) in a flame-dried Schlenk flask under argon. Sodium hydride (NaH; Sigma-Aldrich) was added at a threefold molar excess relative to PEG hydroxyl groups and reacted for 45 min at room temperature. Ethyl 2-(bromomethyl)acrylate (EBrMA; Ambeed, Inc.) was added dropwise at a sixfold molar excess relative to PEG hydroxyl groups, and the reaction proceeded for 48 h at room temperature protected from light. The reaction was quenched with 1 N acetic acid until gas evolution ceased, filtered through Celite 545, and concentrated by rotary evaporation at 50 °C. PEGαMA was precipitated in cold diethyl ether at 4 °C overnight, dried under vacuum, and purified by dialysis against deionized water (1-kg/mol molecular-weight-cutoff membrane) for 4 days. Purified polymer was frozen at −80 °C and lyophilized.

Functionalization was confirmed by proton nuclear magnetic resonance spectroscopy (^1^H NMR). ^1^H NMR (300 MHz, CDCl): δ 1.36 (t, 3H), 3.71 (s, 114H), 4.29 (t, 4H), 5.93 (q, 1H), and 6.34 (q, 1H). Only polymer batches exhibiting >85% α-MA functionalization, calculated from the integration of the vinyl resonances relative to the PEG backbone resonance, were used for hydrogel formation.

### 2.9. Human Lung dECM Preparation and Functionalization

Human lung tissue was decellularized using our established detergent-, salt-, and nuclease-based protocol.^53–55^ Briefly, lung tissue was mechanically disrupted and incubated in 0.1% Triton X-100 (6 mL; Fisher BioReagents) for 24 h at 4 °C, followed by 2% sodium deoxycholate (6 mL; Fisher BioReagents) for 24 h at 4 °C. Samples were subsequently treated with 1 N sodium chloride (6 mL; Chem-Impex) and incubated with DNase (30 mg/L) for 2 h at 30 °C. Decellularized tissues were frozen and lyophilized to obtain dry human lung dECM. Residual double-stranded DNA was quantified using a Quant-iT PicoGreen dsDNA Assay Kit (Thermo Fisher Scientific), confirming <50 ng residual DNA per mg dry tissue.

Lyophilized human lung dECM was solubilized in pepsin-HCl (10 mg dECM/mL; Thermo Fisher Scientific) for 48 h at room temperature and subsequently lyophilized. Free primary-amine content was quantified using a ninhydrin assay (Sigma-Aldrich). Solubilized dECM was thiolated using 2-iminothiolane (Traut’s reagent) in PBS containing 3 mM ethylenediaminetetraacetic acid (EDTA) at pH 7.4. Traut’s reagent was added at a 75-fold molar excess relative to measured primary amines, and the reaction proceeded for 1 h at room temperature. Thiolated dECM (dECM-SH) was purified by dialysis against deionized water using a 0.1-0.5 kg/mol molecular-weight-cutoff membrane (Repligen) and lyophilized. Free-thiol content was quantified using Ellman’s reagent [5,5′-dithiobis-(2-nitrobenzoic acid); Sigma-Aldrich], with absorbance measured using a BioTek Synergy H1 multimode microplate reader. Batch-specific thiol measurements determined the amount of dECM-SH incorporated into each hydrogel formulation.

### 2.10. Hybrid-Hydrogel Fabrication

Hybrid-hydrogels were prepared using eight-arm PEGαMA, thiolated human lung dECM (dECM-SH), and a matrix metalloproteinase-9 (MMP9)-degradable peptide crosslinker (CGGPQGIWGQGC). Formulations contained equal contributions of thiol equivalents from dECM-SH (50%) and the MMP9-degradable peptide crosslinker (50%). Donor-derived dECM-SH was reduced with tris(2-carboxyethyl)phosphine (TCEP; pH 7.6) for 30 min before hydrogel formation. Batch-specific free-thiol concentrations, quantified by Ellman’s assay, determined the amount of dECM-SH and MMP9-degradable peptide crosslinkers incorporated into each formulation.

PEGαMA concentrations were adjusted to achieve tissue-informed, sex- and disease-specific mechanical design criteria derived from female and male control and IPF human lung tissue. PEGαMA concentration was varied independently of dECM donor identity to establish the targeted mechanical conditions. This formulation strategy enabled the incorporation of sex- and disease-specific ECM while tuning network stiffness to reproduce the mechanical relationships measured in human lung tissue. Final formulations are summarized in Table S2.

PEGαMA was equilibrated in Hank’s balanced salt solution (HBSS; pH 7.6) for 5 min before combining with dECM-SH, MMP9-degradable peptide crosslinker, and lithium phenyl-2,4,6-trimethylbenzoylphosphinate (LAP). LAP was incorporated into the precursor formulation before casting, with its solution volume included in the final formulation volume. Precursor solutions were cast between Sigmacote-treated glass slides separated by rubber spacers and exposed to 365-nm light (10 mW/cm²; OmniCure, Lumen Dynamics) for 5 min to initiate LAP-mediated radical photopolymerization. Hydrogels were subsequently transferred to PBS for storage prior to experiments.

Where indicated, secondary crosslinking by dityrosine dimerization between dECM fragments was induced by incubation in a 1:100 ruthenium/sodium persulfate (Ru/SPS) solution for 2–4 h, followed by incubation in fresh PBS for 5 min and exposure to 440-nm light for 5 min. This secondary photochemical reaction provided an additional method of increasing hydrogel stiffness after initial network formation. Hydrogels were characterized under the specified formulation and crosslinking conditions to assess whether the engineered materials reproduced the target human lung mechanical properties.

### 2.11. Rheological Characterization of Hybrid-Hydrogels

Hybrid-hydrogel mechanical properties were measured using a Discovery HR-2 rheometer (TA Instruments) equipped with an 8-mm parallel-plate geometry and a Peltier temperature control system maintained at 37°C. Hydrogels were equilibrated in PBS before measurement. The geometry was lowered until an axial force of 0.03 N was achieved, and the corresponding gap distance was recorded. The gap was subsequently adjusted to achieve 30% compression, corresponding to the compression level at which the storage modulus (G′) reached a plateau. Storage modulus (G′) and loss modulus (G″) were measured, and elastic modulus (E) was calculated using the linear elastic relationship E = 2G′(1 + ν), where ν represents the experimentally determined Poisson’s ratio for each hydrogel formulation (Table S3). Repeated rheological measurements obtained from the same donor-derived hydrogel formulation were considered technical replicates. Donor-level mechanical summaries were used for graphical presentation, with statistical analyses performed as described below.

### 2.12. Immunofluorescence Staining and Analysis of Hybrid-Hydrogels

Acellular hybrid-hydrogels containing female and male control or IPF human lung dECM were stained for COL1A1, tenascin-C (TNC), and periostin (POSTN). Hydrogels were rinsed in PBS and blocked in 5% bovine serum albumin (BSA) in PBS w/v for 2 hours at room temperature. Samples were incubated with primary antibodies diluted in blocking buffer (5% w/v BSA in PBS) overnight at 4 °C. Following three washes in PBS containing 1% Tween-20 (PBST), hydrogels were incubated with species-appropriate fluorescent secondary antibodies overnight at 4 °C, protected from light. Samples were washed in PBS before imaging using a Keyence microscope. Primary and secondary antibodies are listed in Table S4.

Fluorescence images were analyzed using Fiji/ImageJ. Marker-positive area was quantified within the hydrogel ROI and normalized to total hydrogel area. Multiple images acquired from the same donor-derived hydrogel were considered technical or spatial subsamples, with the dECM donor serving as the biological replicate. Results were expressed as the mean percentage of marker-positive area.

### 2.13. Second Harmonic Generation Imaging and Analysis of Human Lung Tissue and Hybrid-Hydrogels

Fibrillar collagen architecture in human lung tissue and hybrid-hydrogels was evaluated using second harmonic generation (SHG) microscopy on a Zeiss LSM780 inverted laser-scanning microscope equipped with a Coherent Chameleon Ultra II pulsed femtosecond laser for two-photon excitation at the University of Colorado Anschutz Advanced Light Microscopy Core. Human lung cryosections (10-µm thickness) were imaged at 10× using 7% laser power and a gain of 650 with six regions were acquired per tissue section. Hybrid-hydrogels were imaged at 10× using 10% laser power and a detector gain of 700. Three regions were acquired per hydrogel as 475-µm z-stacks with 35-µm intervals through the approximately 500-µm hydrogel thickness.

SHG-positive area was quantified as the primary measure of fibrillar collagen architecture, with integrated density evaluated as a complementary measurement. Multiple images from the same tissue or donor-derived dECM were considered spatial subsamples, with the tissue or dECM donor serving as the biological replicate.

### 2.14. Growth Factor Array Analysis of Human Lung dECM

Growth factors retained in human lung dECM were evaluated using a membrane-based human growth factor antibody array (Abcam, ab134002) containing 41 targets, according to the manufacturer’s protocol. To ensure consistent dECM loading across experimental conditions, all donor-derived dECM preparations were resuspended at an equivalent mass concentration before array incubation. Solubilized and thiolated dECM preparations from female and male control and IPF human lungs were loaded at equivalent volumes and mixed 1:1 with blocking buffer to maintain consistent dECM mass per membrane and applied to preblocked membranes overnight at 4 °C. Membranes were washed and sequentially incubated with biotinylated detection antibodies and horseradish peroxidase (HRP)-conjugated streptavidin, with each incubation performed overnight at 4 °C. Following a final wash, membranes were developed by chemiluminescence and imaged using a C-DiGit Blot Scanner (LI-COR). Spot intensities were quantified by densitometry using Fiji/ImageJ, with background subtraction based on negative-control spots and normalization to positive-control spots. The TGF-β1 dataset included 12 donor-derived dECM preparations, with three donors per sex-by-disease condition. Background-corrected measurements were retained as recorded, including genuine zero-valued signals. Array-derived measurements were interpreted as relative TGF-β1 immunoreactivity rather than absolute growth-factor concentration or evidence of biological activity. Multiple measurements from the same dECM preparation were considered technical replicates, with the dECM donor serving as the biological replicate.

### 2.15. Glycosaminoglycan Quantification of Human Lung dECM

Total GAG content in female and male control and IPF human lung dECM was quantified using a Total Glycosaminoglycans Assay Kit (Colorimetric; Abcam, ab289842) according to the manufacturer’s instructions. dECM samples were prepared at 20 mg/mL and incubated with the assay reagents. Absorbance was measured at 400 nm, and GAG concentrations were calculated from a standard curve. GAG content was normalized to dECM mass. Technical replicate measurements were summarized for each dECM preparation, with the donor serving as the biological replicate.

### 2.16. Alcian Blue Staining and Analysis of Hybrid-Hydrogels

Hybrid-hydrogels were stained with Alcian Blue to evaluate acidic GAG-rich matrix retained following dECM processing and hydrogel fabrication. Whole-mount hydrogels were handled using a modified staining procedure to accommodate the greater thickness and hydrated structure of the engineered materials.

Hydrogels were fixed in 4% paraformaldehyde for 30 min, washed three times in PBS for 8–10 min per wash, and equilibrated in 3% acetic acid for 5 min. Samples were incubated in 1% (w/v) Alcian Blue 8GX prepared in 3% acetic acid (pH 2.5) for 60 min at room temperature. Following staining, hydrogels were washed twice in 3% acetic acid for 5 min per wash and three times in deionized water for 10 min per wash. Sample transfers and washes were performed gently to minimize hydrogel deformation or loss of material.

Unlike cryosectioned human lung tissue, whole-mount hydrogels were maintained in a hydrated state for imaging rather than subjected to ethanol dehydration and clearing. Brightfield images were acquired under consistent settings across experimental groups. Images were analyzed in Fiji/ImageJ, with ROIs defined to exclude surrounding background. Alcian Blue-positive area was quantified using a consistent color-thresholding procedure and normalized to total hydrogel area. The percentage of Alcian Blue-positive area was calculated as 100 × (Alcian Blue-positive area / total hydrogel ROI area).

The finalized hybrid-hydrogel Alcian Blue analysis included 121 images from 36 hydrogels representing 12 donor-derived ECM preparations, with three biological donors per sex-by-disease condition. Multiple images from the same donor-derived hybrid-hydrogel were treated as technical or spatial subsamples, with the dECM donor serving as the biological replicate. Statistical models accounted for image measurements nested within individual hydrogels and hydrogels nested within donor-derived ECM preparations.

### 2.17. Comparison of Human Lung Tissue and Hybrid-Hydrogel Properties

Fidelity between human lung tissue and hybrid-hydrogels was evaluated across four prespecified biological contrasts: female IPF relative to female control, male IPF relative to male control, female control relative to male control, and female IPF relative to male IPF. Technical and spatial measurements were summarized at the donor level using the assay-specific procedures described above. For positive-valued endpoints, condition-level effects were expressed as log_2_ fold changes.

Direct human lung tissue-to-hybrid-hydrogel comparisons were performed for SHG, COL1A1 immunofluorescence, TNC immunofluorescence, POSTN immunofluorescence, and Alcian Blue staining. Human lung tissue AFM and hybrid-hydrogel rheology were evaluated as complementary measurements of matrix mechanics. Comparisons between human lung tissue structural-proteoglycan abundance and processed-dECM total GAG concentration, and between human lung tissue TGF-β1 proteomic abundance and TGF-β1 immunoreactivity in processed dECM, were considered biologically related but analytically non-equivalent measurements.

Human lung tissue-to-hybrid-hydrogel effect sizes were plotted relative to the line of identity (y = x). Lin’s concordance correlation coefficient (CCC) was calculated to assess absolute agreement between analytically comparable effects, and Spearman’s rank correlation coefficient (ρ) was calculated to evaluate preservation of effect ordering. Because each analysis contained four prespecified biological contrasts rather than four independent experimental samples, concordance and correlation statistics were interpreted descriptively rather than as independent n = 4 hypothesis tests.

### 2.18. Statistical Analysis

The human donor, or the donor from which a dECM preparation was derived, was considered the biological experimental unit throughout the study. Images, ROIs, AFM indentations, repeated rheological measurements, replicate assay wells, and repeated antibody-array measurements were considered spatial or technical subsamples and were not treated as independent biological replicates.

For datasets containing repeated measurements within each donor, linear mixed-effects models were fitted by restricted maximum likelihood (REML) in JMP. Disease status, sex, and the disease-by-sex interaction were included as fixed effects, with donor included as a random effect. Response variables exhibiting substantial right skew were log10-transformed prior to modeling.

For human lung AFM, SHG, immunofluorescence, and Alcian Blue datasets, repeated measurements were modeled with donor as the biological grouping variable. For hybrid-hydrogel SHG and Alcian Blue analyses, gel identity was nested within donor where the underlying data included individual gel identifiers. For hydrogel immunofluorescence, donor-level random effects were used to account for repeated image measurements because gel identifiers were not consistently available in the analyzed dataset.

For assays yielding one summarized value per donor or dECM source, including total GAG and antibody-array measurements, donor-level values were analyzed using a two-factor linear model containing disease status, sex, and the disease-by-sex interaction. Four pairwise comparisons were specified a priori: female IPF versus female control, male IPF versus male control, female control versus male control, and female IPF versus male IPF. Pairwise estimates were obtained from interaction-group least-squares means with Holm adjustment across the four prespecified comparisons. Overall fixed-effect tests were evaluated for disease status, sex, and the disease-by-sex interaction. Where applicable, both the overall model effects and corresponding planned pairwise comparisons were reported to distinguish broad disease- or sex-associated effects from differences between specific experimental groups.

GraphPad Prism was used for graphical presentation. Values imported into Prism represented donor-level summaries unless otherwise indicated; consequently, each plotted symbol represents an independent donor or donor-derived source rather than an individual technical measurement. Exact biological sample sizes, transformations, error-bar definitions, and statistical models are reported in the corresponding figure legends.

Statistical significance was defined as p < 0.05 unless otherwise specified. Values with 0.05 ≤ p ≤ 0.10 were described as trends, consistent with the reporting convention used throughout this manuscript. Statistical interpretations were based on unrounded p-values. Holm-adjusted p-values were used for the prespecified pairwise comparisons presented in the main figures, with unadjusted values reported separately where relevant.

Correspondence between human lung tissue and hybrid-hydrogels was evaluated across four prespecified biological contrasts: female IPF relative to female control, male IPF relative to male control, female control relative to male control, and female IPF relative to male IPF. Technical and spatial measurements were summarized at the donor level using the assay-specific procedures described above. For positive-valued endpoints, condition-level effects were expressed as log□ fold changes.

Direct human lung tissue-to-hybrid-hydrogel comparisons were performed for SHG, COL1A1 immunofluorescence, TNC immunofluorescence, POSTN immunofluorescence, and Alcian Blue staining. Human lung tissue AFM and hybrid-hydrogel rheology were evaluated as complementary measurements of matrix mechanics.

Comparisons between human lung tissue structural-proteoglycan abundance and processed-dECM total GAG content were treated as biologically related but analytically non-equivalent measurements. Native-lung TGF-β1 proteomic abundance and TGF-β1 signal retained in processed dECM were also considered complementary measurements. The native-lung TGF-β1 proteomic comparison involved control alveolar tissue and IPF fibroblastic-focus regions, whereas the processed-dECM measurement represented material recovered following tissue processing.

Human lung tissue-to-hybrid-hydrogel effect sizes were plotted relative to the line of identity (y = x). Lin’s concordance correlation coefficient (CCC) was calculated to assess absolute agreement between analytically comparable effects, and Spearman’s rank correlation coefficient (ρ) was calculated to evaluate preservation of effect ordering. Pearson’s correlation coefficient (r) was additionally used to describe the linear association between corresponding tissue and material effect estimates.

## 3. Results and Discussion

### 3.1. Spatial proteomics reveals region- and sex-associated differences in human lung matrisome

To define human lung ECM features that inform biomaterial design, we performed a matrisome-focused analysis of a previously published region-resolved LCM-MS proteomic dataset comprising control alveoli, morphologically normal alveoli from IPF lungs (IPF alveoli), and fibroblastic foci from female and male donors (Fig 1A).^33, 34^ Hierarchical clustering of matrisome proteins revealed distinct abundance patterns across anatomical regions, with additional variation associated with donor sex (Fig. 1B). Fibroblastic foci exhibited the greatest divergence from alveolar regions, consistent with localized ECM remodeling in IPF. Previous spatial characterization of human IPF lung similarly identified regional variation in collagen, glycoprotein, and proteoglycan distribution within fibroblastic foci.^32^ Region- and disease-associated differences in ECM composition have also been reported in decellularized human lung tissue.^36, 48^ Several proteins identified within the overview heatmap were particularly relevant to subsequent characterization of the fibrotic ECM (Fig. 1B). These included the matricellular glycoproteins tenascin-C (TNC) and periostin (POSTN), which contribute to cell–matrix interactions and fibrotic remodeling,^65, 66^ and transforming growth factor-β-induced protein (TGFBI), an ECM glycoprotein associated with TGF-β-responsive signaling and pulmonary fibrosis.^67^ The small leucine-rich proteoglycans lumican (LUM) and decorin (DCN) were also identified and are important regulators of collagen fibrillogenesis and matrix organization.^68^ These matrix components contribute to tissue structure, cell-matrix signaling, and ECM organization, highlighting the importance of characterizing fibrotic ECM beyond collagen abundance alone. Together, these findings motivated subsequent evaluation of tissue mechanics, fibrillar collagen organization, matricellular proteins, and GAG-rich matrix in female and male human lung tissue.^15, 32, 40^

**Figure 1.**
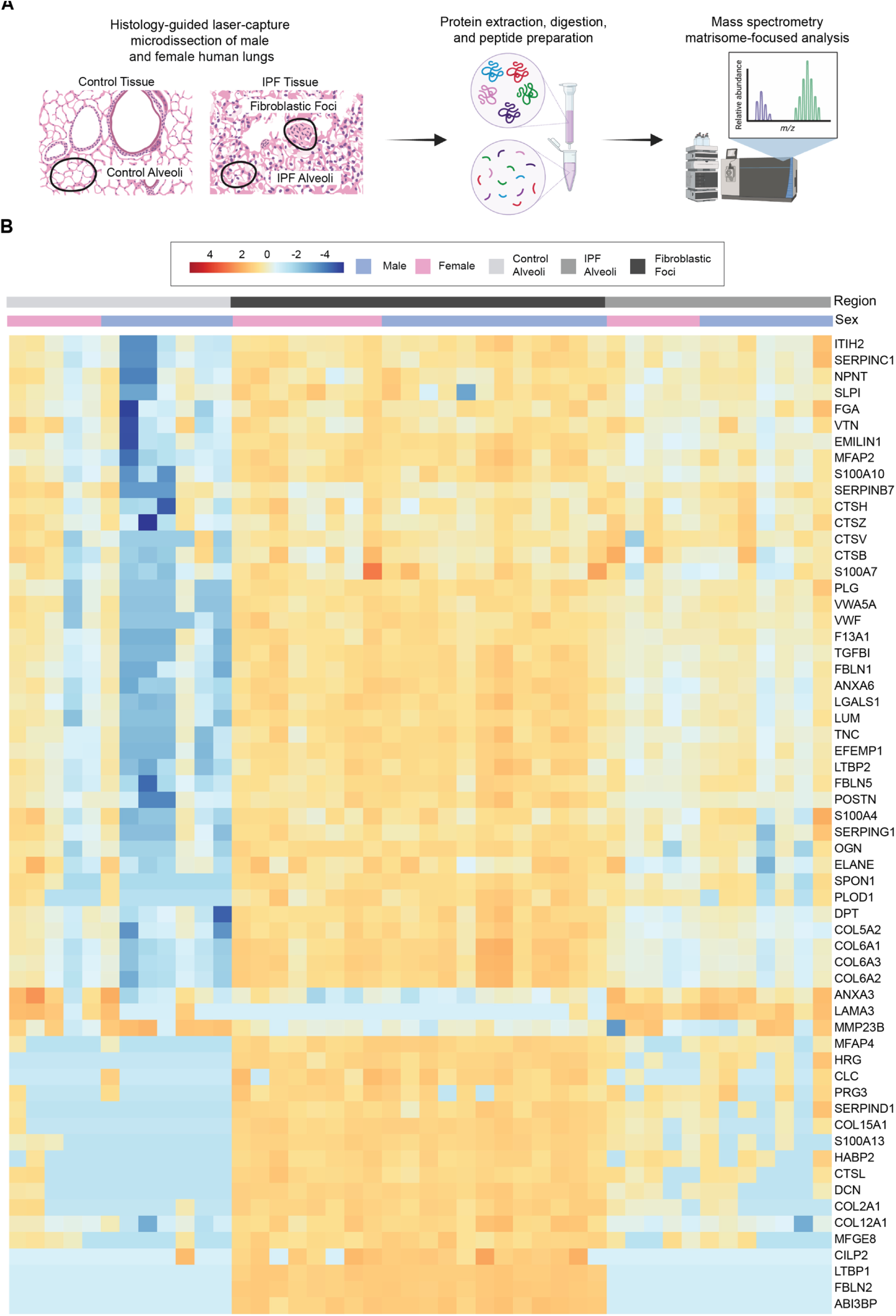
Spatial matrisome proteomics reveals region- and sex-associated differences in human lung ECM. (A) Experimental workflow for histology-guided laser capture microdissection of control alveoli, IPF alveoli, and fibroblastic foci from female and male human lung tissue, followed by protein extraction, digestion, mass spectrometry, and matrisome-focused analysis. (B) Hierarchically clustered heatmap of matrisome protein abundance across individual samples. Selected proteins include the matricellular glycoproteins TNC and POSTN, the TGF-β-inducible matrix protein TGFBI, and the small leucine-rich proteoglycans LUM and DCN. Annotation bars indicate anatomical region and donor sex, and the heatmap displays row-scaled protein abundance.

### 3.2. Female and male lungs exhibit different regional matrisome remodeling programs

We next analyzed female and male matrisomes separately to determine whether IPF-associated ECM remodeling involved similar molecular programs in both sexes. Within each sex, hierarchical clustering revealed changes in protein abundance across control alveoli, IPF alveoli, and fibroblastic foci (Fig. 2A–B). These regional patterns included variation in collagens (COL1A1, COL5A2, and COL6A3), ECM glycoproteins (POSTN and FN1), and proteoglycans (DCN, BGN, VCAN, and HSPG2). Notably, COL1A1 was represented in both sex-stratified heatmaps, whereas POSTN appeared among the selected proteins displayed in the female heatmap (Fig. 2A–B). Rather than uniform accumulation of ECM proteins, the heatmaps revealed differences in the relative abundance of matrix components across regions representing different pathological states of IPF. Previous studies have similarly identified regional differences in human lung ECM composition and characterized fibroblastic foci as sites of active collagen biosynthesis.^36, 69^ Notably, morphologically normal IPF alveoli exhibited molecular differences from control alveoli, suggesting that ECM remodeling may occur in regions without fibrosis identifiable by histology.

**Figure 2.**
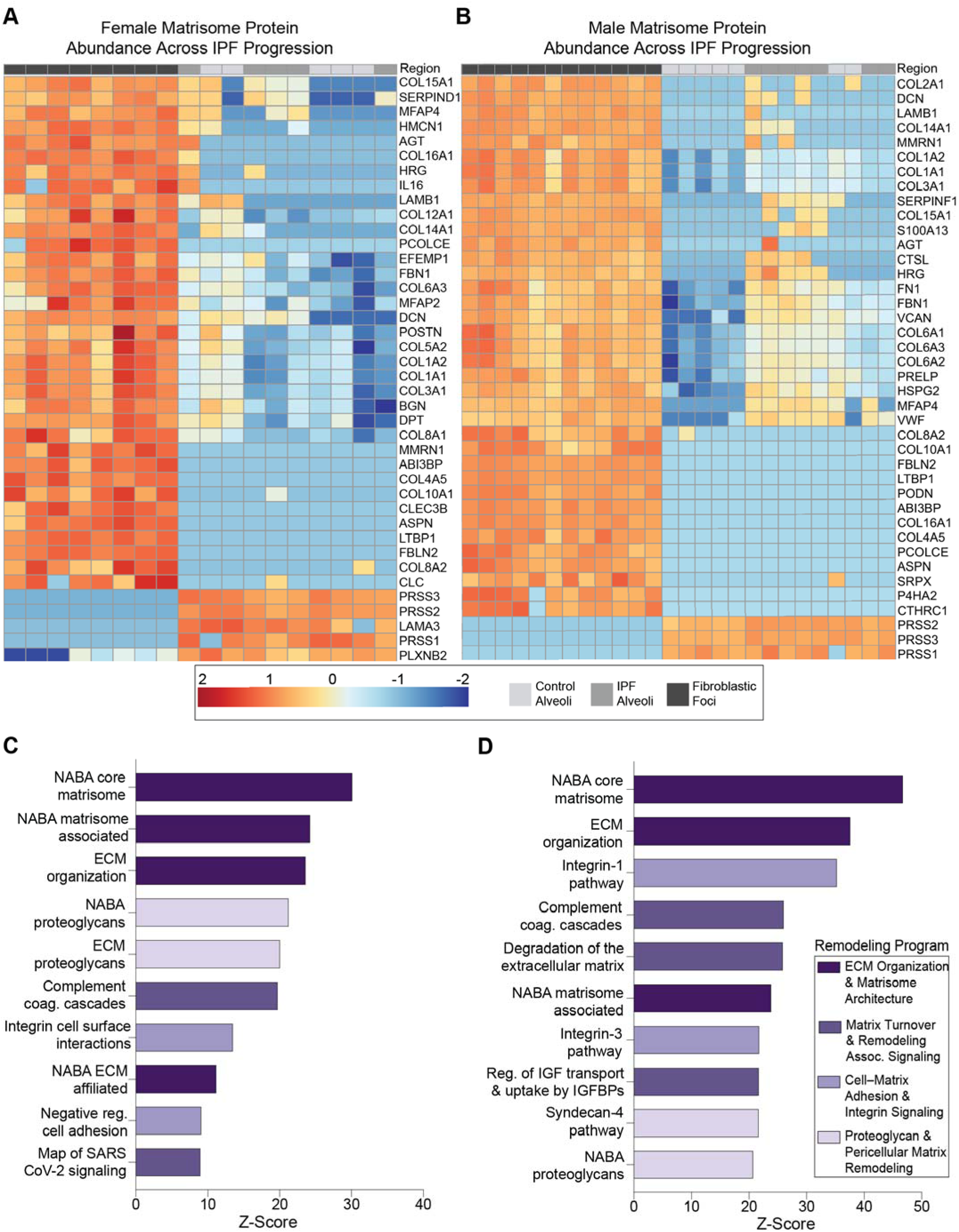
Female and male lungs exhibit distinct regional matrisome remodeling patterns and overlapping functional remodeling programs. Hierarchically clustered heatmaps of selected matrisome proteins analyzed separately in (A) female and (B) male control alveoli, IPF alveoli, and fibroblastic foci. Annotation bars identify anatomical region, and heatmap color represents row-scaled protein abundance. Selected proteins include COL1A1, which is represented in both sex-stratified heatmaps, and POSTN, which is displayed in the female heatmap. Pathway enrichment analysis of matrisome proteins in (C) female and (D) male lungs. Enriched pathways were organized by biological function into four remodeling programs: ECM organization and matrisome composition, matrix turnover and remodeling-associated signaling, cell–matrix adhesion and integrin signaling, and proteoglycan/pericellular-matrix remodeling. These functional groupings were assigned based on established matrisome classifications and shared biological functions rather than an additional statistical clustering analysis.

Pathway enrichment further identified overlapping but nonidentical ECM-associated pathways and gene sets in female and male lungs (Fig. 2C–D). To interpret these enriched pathways, we organized them into four functional remodeling categories based on established matrisome classifications and the biological processes through which ECM components regulate tissue structure, matrix turnover, and cell–matrix interactions.^30, 61, 62^ These categories were assigned according to shared biological functions to facilitate interpretation of the enrichment results, rather than generated through an additional statistical clustering analysis.

The ECM organization and matrisome composition program included NABA core matrisome, NABA matrisome-associated, NABA ECM-affiliated, and ECM organization terms. These gene sets encompass structural and ECM-associated constituents, including collagens and matrix glycoproteins, that contribute to extracellular matrix composition and organization.^30, 61, 62^ The matrix turnover and remodeling-associated signaling program included ECM degradation, complement/coagulation cascades, and regulation of IGF transport and uptake by IGF-binding proteins. These pathways were grouped based on their involvement in matrix degradation, coagulation-associated fibrotic remodeling, and the regulation of matrix-associated growth-factor availability.^30, 70^

Cell–matrix adhesion and integrin signaling encompassed integrin-1 and integrin-3 pathways, integrin cell-surface interactions, and negative regulation of cell adhesion. These pathways regulate interactions between ECM ligands and cell-surface receptors and are relevant to fibroblast adhesion, mechanosensing, and profibrotic signaling.^63^ Finally, proteoglycan and pericellular matrix remodeling included NABA proteoglycans, ECM proteoglycans, and syndecan-4 signaling. This grouping reflects the roles of proteoglycans and their associated GAG chains in collagen organization, pericellular matrix interactions, and growth-factor regulation.^64, 68, 71^

Both datasets were enriched for core-matrisome and ECM-organization pathways. Female samples additionally exhibited enrichment of NABA proteoglycans, ECM proteoglycans, integrin cell-surface interactions, and negative regulation of cell adhesion. Male samples showed enrichment of integrin-1 and integrin-3 pathways, ECM degradation, regulation of IGF transport by IGF-binding proteins, and syndecan-4 signaling. Complement/coagulation cascades and proteoglycan-associated pathways were represented in both datasets. These differences suggest that sex-associated ECM remodeling may involve variation in matrix organization, turnover, and cell–matrix communication rather than structural protein abundance alone. Importantly, the results indicate differences in the particular pathways represented within shared remodeling functions, rather than entirely sex-exclusive ECM remodeling mechanisms.

The enrichment of integrin-associated pathways in male lungs is relevant to matrix mechanosensing and fibroblast activation,^16^ while proteoglycan-associated pathways in female lungs suggest differences in matrix hydration and growth-factor presentation. These findings are consistent with established features of pulmonary fibrosis. Previous studies have identified collagen biosynthesis, ECM degradation, and integrin-associated remodeling within fibroblastic foci, while investigations of human IPF tissue have also reported increased GAG deposition and altered heparan sulfate structure.^32, 69, 71^ Thus, both the integrin- and turnover-associated signatures represented in male lungs and the proteoglycan-associated signatures represented in female lungs overlap with previously described IPF biology. The present sex-stratified analyses suggest that the relative representation of these processes may differ between female and male lung ECM.

This distinction is particularly relevant because IPF is more prevalent in males, and clinical IPF cohorts frequently include substantially more male than female patients.^21^ Consequently, descriptions of a typical fibrotic ECM phenotype may not fully capture remodeling patterns present in female IPF lungs, although male-predominant clinical sampling does not establish that previously reported molecular signatures are specifically male-derived. Previous characterization of decellularized human lung ECM demonstrated region- and disease-associated changes in GAG composition and function, supporting the importance of this matrix component in pulmonary disease.^48^ Proteoglycans and associated GAG chains regulate collagen organization, growth-factor availability, and cell–matrix signaling through differences in sulfation, charge density, hydration, and ligand-binding properties.^48, 51, 64, 68, 71^ The enrichment of proteoglycan- and pericellular-matrix-associated pathways in female lungs highlights key biochemical and structural aspects of ECM remodeling that may not be captured by collagen abundance or tissue stiffness alone.

Together, these pathway-level differences suggest that female and male IPF lungs may differ in the relative contributions of structural matrix composition, matrix turnover, cell–matrix signaling, and proteoglycan-associated remodeling. Because pathway enrichment was performed separately within the female and male datasets, these patterns should be interpreted as descriptive differences in pathway representation rather than direct statistical evidence of sex-dependent enrichment. These differences extend the cell-adhesion and ECM-organization programs previously identified in distal IPF regions and highlight the limitations of defining fibrotic microenvironments and biomaterial models by elevated collagen content or stiffness alone.^12, 34, 45, 46^ Accordingly, we focused subsequent analyses on four complementary matrix properties: matrix mechanics, fibrillar collagen organization, matricellular proteins, and GAG-rich matrix. These findings established the foundation for sex- and disease-informed biomaterial design criteria derived directly from human lung tissue by integrating complementary measurements of matrix composition, organization, and mechanics.

### 3.3. Sex- and region-associated remodeling alters core-matrisome abundance and composition

Analysis of total normalized matrisome protein intensity revealed substantial regional variation, with fibroblastic foci exhibiting greater matrisome signal than alveolar regions in both sexes and the highest signal observed in male fibroblastic foci (Fig. 3A). However, these patterns were not uniform across disease states, as the relative female-male relationship differed between control alveoli and IPF regions. Changes in total matrisome signal were accompanied by shifts in core-matrisome composition. Although, collagens represented the largest fraction across conditions, IPF alveoli contained proportionally greater contributions from ECM glycoproteins and proteoglycans than fibroblastic foci (Fig. 3B). This redistribution was particularly apparent in male IPF alveoli, where glycoproteins accounted for a larger proportion of the matrisome relative to collagen.

**Figure 3.**
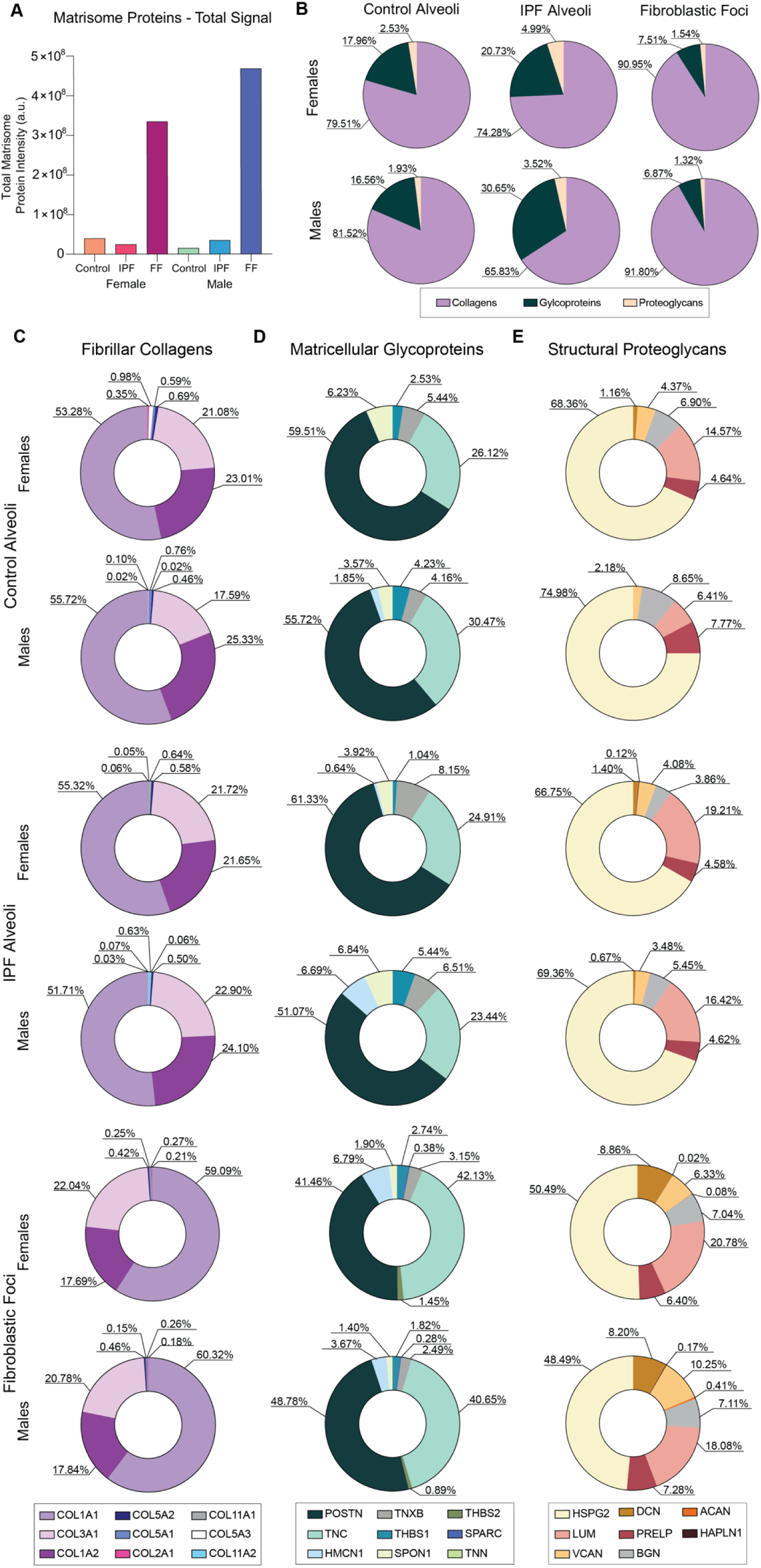
Sex-associated ECM remodeling reveals distinct regional changes in core-matrisome composition. (A) Total normalized matrisome protein intensity across female and male control alveoli, IPF alveoli, and fibroblastic foci. (B) Relative contributions of collagens, ECM glycoproteins, and proteoglycans to core matrisome within each sex and anatomical region. Relative contributions of individual proteins within the (C) fibrillar collagen, (D) matricellular glycoprotein, and (E) structural proteoglycan groups, expressed as percentages of the corresponding protein group.

Examination of individual protein contributions within the fibrillar collagen (Fig. 3C), matricellular glycoprotein (Fig. 3D), and structural proteoglycan (Fig. 3E) groups revealed additional compositional heterogeneity across female and male lung regions. The relative contributions of individual proteins shifted across control, IPF alveoli, and fibroblastic foci, with different distributions observed between sexes. These findings reinforce that ECM remodeling involves changes in both total matrisome protein intensity and the molecular composition of individual matrix classes.

Together, these analyses reveal distinct sex-associated patterns of ECM remodeling across anatomical regions representing different pathological states of IPF. Rather than simply differing in the magnitude of shared fibrotic response, female and male lungs exhibited different shifts in total matrisome signal, core-matrisome composition, and individual protein contributions. Although these regions do not represent a temporal sequence, the molecular profiles are consistent with divergent ECM remodeling trajectories across disease-associated states. Sex-associated differences in ECM composition have also been identified in cartilage and cardiac tissues, indicating that divergent matrix remodeling may represent a broader feature of tissue homeostasis and disease.^13, 14^ These differences establish the compositional foundation for sex- and disease-informed biomaterial design criteria, motivating subsequent characterization of human lung mechanics, fibrillar collagen organization, matricellular proteins, and GAG-rich matrix, followed by evaluation of the preservation of these tissue-derived properties in engineered hybrid-hydrogels.

### 3.4. Sex-stratified human lung mechanics reveal distinct fibrotic remodeling patterns recapitulated in hybrid-hydrogels

We next investigated whether sex- and disease-associated mechanical properties of human lung tissue could be recapitulated in engineered hybrid-hydrogels. Donor-derived lung dECM was incorporated into PEGαMA hybrid-hydrogels using complementary crosslinking mechanisms: LAP-mediated radical polymerization established the initial network, while Ru/SPS-mediated dityrosine crosslinking enabled secondary stiffening (Fig. 4A). Hybrid-hydrogel formulations were tailored to reproduce the mechanical properties measured in female and male control and IPF human lung tissue, with male IPF hydrogels fabricated either at IPF-rang stiffness or initially at control-range stiffness for subsequent dynamic stiffening during culture. Unlike conventional ECM-derived hydrogels, in which composition and mechanics are often coupled,^46, 50^ this hybrid-hydrogel approach combines donor-specific biochemical cues with independent control over matrix stiffness.^34, 54, 55, 72^ To evaluate the fidelity of these engineered microenvironments, we compared human lung tissue and hybrid-hydrogels across four tissue-informed design criteria: matrix mechanics, fibrillar collagen organization, matricellular glycoproteins, and proteoglycan/GAG-rich matrix (Table 1).

**Figure 4.**
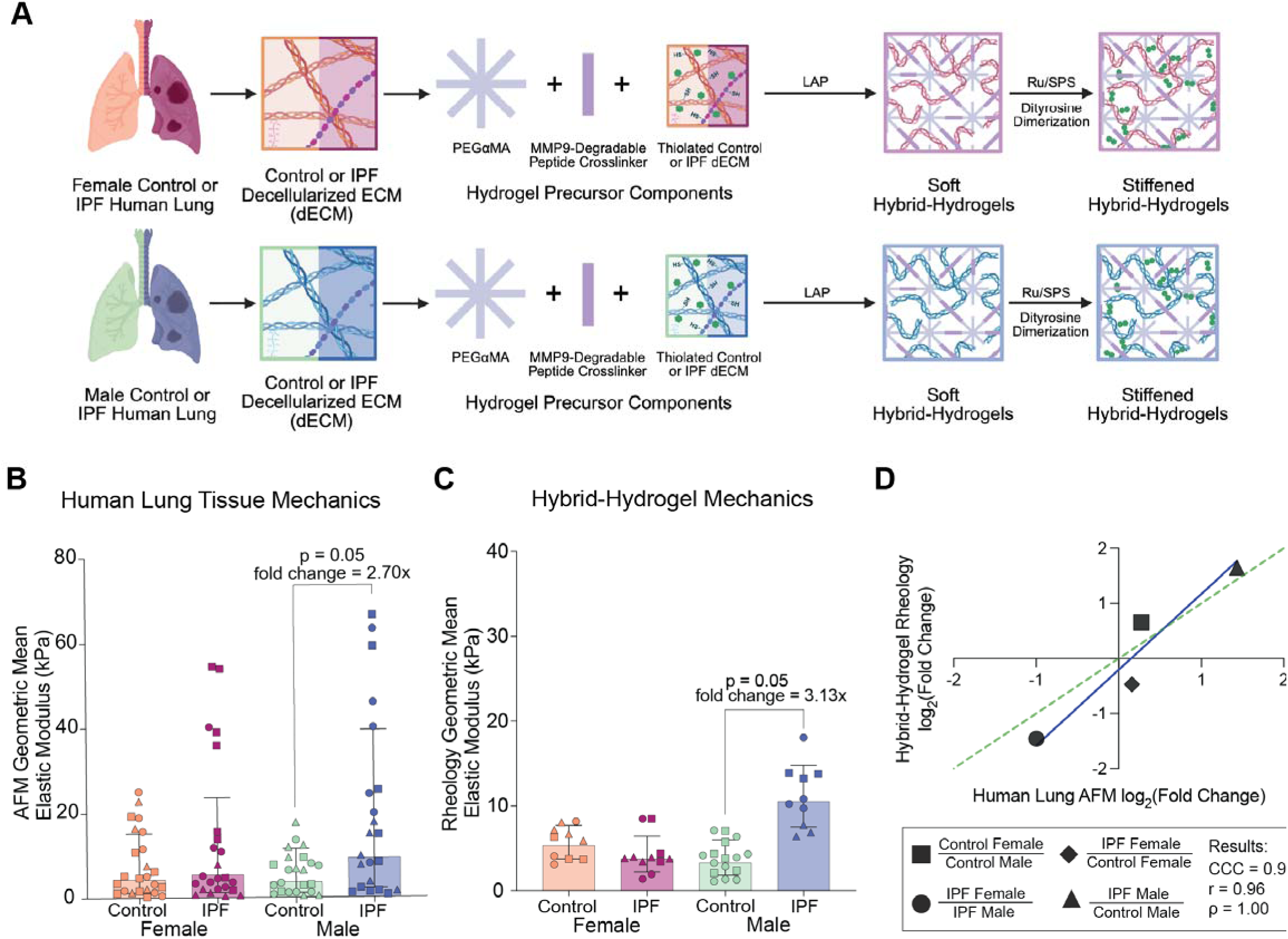
Sex-stratified human lung mechanics reveal distinct fibrotic remodeling patterns recapitulated in hybrid-hydrogels. (A) Fabrication of female and male control and IPF hybrid-hydrogels incorporating donor-derived human lung dECM into PEGαMA networks with an MMP9-degradable peptide crosslinker. LAP-mediated photopolymerization established the initial hydrogel network, while Ru/SPS-mediated dityrosine crosslinking enabled secondary stiffening. (B) Geometric mean elastic modulus of female and male control and IPF human lung tissue measured by AFM. Male IPF tissue exhibited a 2.70-fold increase in elastic modulus relative to male control tissue (p=0.050). (C) Geometric mean elastic modulus of corresponding hybrid-hydrogels measured by rheology. Male IPF-derived hybrid-hydrogels exhibited a 3.13-fold increase in elastic modulus relative to male control-derived hybrid-hydrogels (p=0.050). (D) Human lung-to-hybrid-hydrogel mechanical effect comparison across four predefined sex- and disease-associated contrasts, expressed as log_2_ fold changes. The dashed line represents the line of identify. Mechanical effects exhibited strong descriptive concordance (Lin’s concordance correlation coefficient (CCC) = 0.91; r=0.96), supporting successful translation of tissue-informed mechanical design criteria into engineered hybrid-hydrogels.

**Table 1.** Tissue-informed design criteria and corresponding measurements used to evaluate hybrid-hydrogel fidelity. Four ECM dimensions identified through human lung characterization were evaluated using complementary measurements in female and male control and IPF human lung tissue and donor-derived hybrid-hydrogels. These analyses assess the preservation of mechanical, structural, and biochemical matrix features following dECM processing and hydrogel fabrication.

| Tissue Informed | Human Lung Measurement | dECM Hybrid-Hydrogel Measurement |
| --- | --- | --- |
| Lung Mechanics | Atomic Force Microscopy | Rheology |
| Fibrillar Collagen | Second Harmonic Generation<br>Immunofluorescence | Second Harmonic Generation<br>Immunofluorescence |
| Matricellular Glycoproteins | LCM-MS Proteomics<br>Immunofluorescence | Immunofluorescence |
| Proteoglycans/GAG Matrix | LCM-MS Proteomics<br>Alcian Blue Staining | Alcian Blue Staining |

Our sex-stratified AFM analysis of human control and IPF lung tissue revealed pronounced sex-associated differences in disease-associated mechanical remodeling between females and males (Fig. 4B). Male IPF tissue exhibited a 2.70-fold increase in elastic modulus relative to male control tissue (p=0.050), whereas female IPF tissue showed no statistically significant disease-associated change (p=0.453). These findings identify different patterns of mechanical remodeling in female and male IPF lungs, challenging the assumption that increased stiffness is a uniform feature of pulmonary fibrosis across biological sex. Importantly, these mechanical patterns occurred despite extensive ECM remodeling in both sexes (Figs. 1-3), indicating that biochemical changes in the fibrotic matrix do not necessarily translate into equivalent changes in tissue mechanics.

These findings extend previous sex-associated differences in experimental pulmonary fibrosis to human lung tissue. Redente et al. reported greater fibrosis-associated stiffening in male than female mice following bleomycin injury,^44^ while our previous studies identified sex- and age-associated differences in fibroblast responses to dynamic matrix stiffening in engineered pulmonary vascular models.^43^ Together, these studies support a broader role for biological sex in pulmonary mechanobiology. The present human data add an important tissue-level dimension by identifying sex-associated mechanical microenvironments that may influence epithelial and fibroblast responses during IPF. This distinction is biologically relevant because increased stiffness can reinforce fibroblast activation through mechanotransduction pathways, including YAP/TAZ signaling, and establish feedback loops that sustain ECM deposition and myofibroblast persistence.^15, 41, 42^

The divergence between ECM composition and tissue stiffness highlights the importance of higher-order matrix organization. Collagen organization, crosslinking, proteoglycan-associated hydration, and glycoprotein-mediated interactions collectively influence tissue mechanics.^16, 35, 37, 40^ Previous proteomic studies have identified changes in collagen isoform composition and post-translational modifications that may contribute to fibrillar organization, crosslinking, and matrix mechanics.^37, 38, 73, 74^ Thus, increased COL1A1 abundance in some female IPF regions^34^ may not translate into greater tissue stiffness if collagen assembly and higher-order fibrillar organization differ from male IPF lung. These findings reinforce that collagen abundance alone cannot adequately predict the mechanical properties of female and male fibrotic lung tissue.

The disease-associated mechanical patterns observed in human lung tissue were recapitulated in donor-derived hybrid-hydrogels (Fig. 4C). Male IPF-derived hydrogels exhibited a 3.13-fold increase in elastic modulus relative to male control-derived hydrogels (p = 0.050), closely matching the 2.70-fold increase measured in human lung tissue. Female-derived hydrogels exhibited comparatively modest disease-associated differences, consistent with the human lung AFM findings. Previous studies have shown that human lung hybrid-hydrogels can reproduce disease-associated mechanical properties, while crosslinking strategies provide additional control over stiffness.^34, 53–55^ Our findings extend these approaches by demonstrating that sex- and disease-informed mechanical design criteria can be translated into a hybrid-hydrogel platform. Importantly, these mechanical properties were intentionally engineered to match the corresponding human lung conditions rather than relying solely on mechanical properties retained through dECM processing.

To quantify the correspondence between human lung tissue and engineered materials, we compared log fold changes in elastic modulus across four predefined sex- and disease-associated contrasts derived from human lung AFM and hybrid-hydrogel rheology measurements (Fig. 4D). These comparisons exhibited strong descriptive concordance between human lung tissue and hybrid-hydrogels (Lin’s concordance correlation coefficient = 0.91; r = 0.96), with effects distributed near the line of identity. Notably, the male IPF versus male control comparison exhibited closely matched disease-associated increases in elastic modulus, with a 2.70-fold increase in human lung tissue and a 3.13-fold increase in hybrid-hydrogels. Because these comparisons represent four predefined biological contrasts rather than independent experimental samples, concordance statistics were interpreted descriptively. Nevertheless, this agreement supports successful translation of sex- and disease-associated mechanical relationships from human lung tissue into engineered biomaterials.

Our results establish that sex- and disease-informed mechanical design criteria can be recapitulated in engineered hybrid-hydrogels. The pronounced mechanical remodeling observed in male IPF lung, contrasted with the comparatively modest disease-associated change in female lung, highlights the limitations of defining a single fibrotic stiffness for both sexes. By reproducing these distinct mechanical environments while maintaining independent control over ECM composition and stiffness, the hybrid-hydrogel platform provides a foundation for investigating how sex-associated matrix properties influence cellular responses. We next evaluated whether additional tissue-derived design criteria, including fibrillar collagen architecture, matricellular proteins, and proteoglycan/GAG-rich matrix, were retained following dECM processing and hybrid-hydrogel fabrication.

### 3.5. Fibrillar collagen organization is altered during hybrid-hydrogel fabrication despite preservation of COL1A1-associated abundance patterns

We next used SHG imaging to examine fibrillar collagen organization and investigate its relationship with sex-associated differences in human lung tissue mechanics. Unlike COL1A1 immunofluorescence, SHG detects organized fibrillar collagen through its non-centrosymmetric structure, providing complementary information about collagen remodeling. Mixed-effects analysis of human lung SHG-positive area identified a significant overall effect of disease status (p = 0.0305) and a trend toward a sex × disease interaction (p = 0.0766). Human IPF lung tissue exhibited greater and more spatially concentrated SHG-positive fibrillar collagen than control tissue, with a trend toward increased SHG-positive area in the male IPF lungs compared with male controls (p=0.06; Fig. 5A). Together, the significant disease-associated increase and the trend toward a sex × disease interaction suggest that the magnitude of fibrillar collagen remodeling may differ between female and male IPF lungs. This pattern paralleled the pronounced disease-associated stiffening observed in male IPF lung, highlighting a potential relationship between fibrillar collagen organization and tissue mechanics. By complementing protein abundance measurements, SHG provides additional insight into the higher-order organization of the fibrotic matrix. This distinction is functionally relevant because collagen organization influences cell behavior: IPF-like collagen organization and alignment alter fibroblast migration, while enzymatic crosslinking by lysyl oxidases contributes to pathological fibrillar remodeling and matrix mechanics.^40, 75^ Integrating SHG with collagen-specific measurements therefore provides a more comprehensive characterization of ECM remodeling and its potential mechanobiological consequences.

SHG-positive fibrillar collagen remained detectable following decellularization, digestion, functionalization, and incorporation into PEGαMA hybrid-hydrogels (Fig. 5C). However, the fibrillar signal was substantially reduced compared with human lung tissue, and the sex- and disease-associated patterns were not consistently preserved. Although donor-derived ECM retained components capable of contributing to SHG-active fibrillar structures, extensive processing and reconstitution altered the higher-order collagen organization observed in human lung tissue. Consistent with these observations, the human lung-to-hybrid-hydrogel SHG comparison exhibited limited positive concordance across the four predefined biological contrasts (CCC=0.31; Fig. 5E), indicating that sex- and disease-associated patterns of fibrillar collagen organization were only partially reproduced in the engineered materials.

COL1A1 immunofluorescence provided a complementary molecular assessment of collagen remodeling (Fig. 5B). Unlike SHG, COL1A1-positive area did not exhibit a comparable disease-associated increase in male IPF human lung tissue. This distinction was also apparent in hybrid-hydrogels, where male IPF-derived materials retained detectable SHG-positive fibrillar structures despite relatively low COL1A1-postive area (Fig. 5D). One possible explanation is that SHG-active structures contain contributions from other fibrillar collagens, including collagen III, while differences in fibril organization, crosslinking, or antibody epitope accessibility may also influence these measurements. Previous proteomic studies identified changes in collagen isoform composition and post-translational modifications in pulmonary fibrosis, supporting the biological plausibility of these mechanisms.^37, 38^ However, the present experiments do not establish which collagen isoforms or structural modifications contribute to the observed differences. Notably, COL1A1-associated effects exhibited substantially stronger human lung-to-hybrid-hydrogel concordance (CCC = 0.88; Fig. 5F) than SHG-defined fibrillar organization (CCC = 0.31; Fig. 5E), indicating that relative COL1A1 abundance patterns were more consistently preserved following hybrid-hydrogel fabrication.

**Figure 5.**
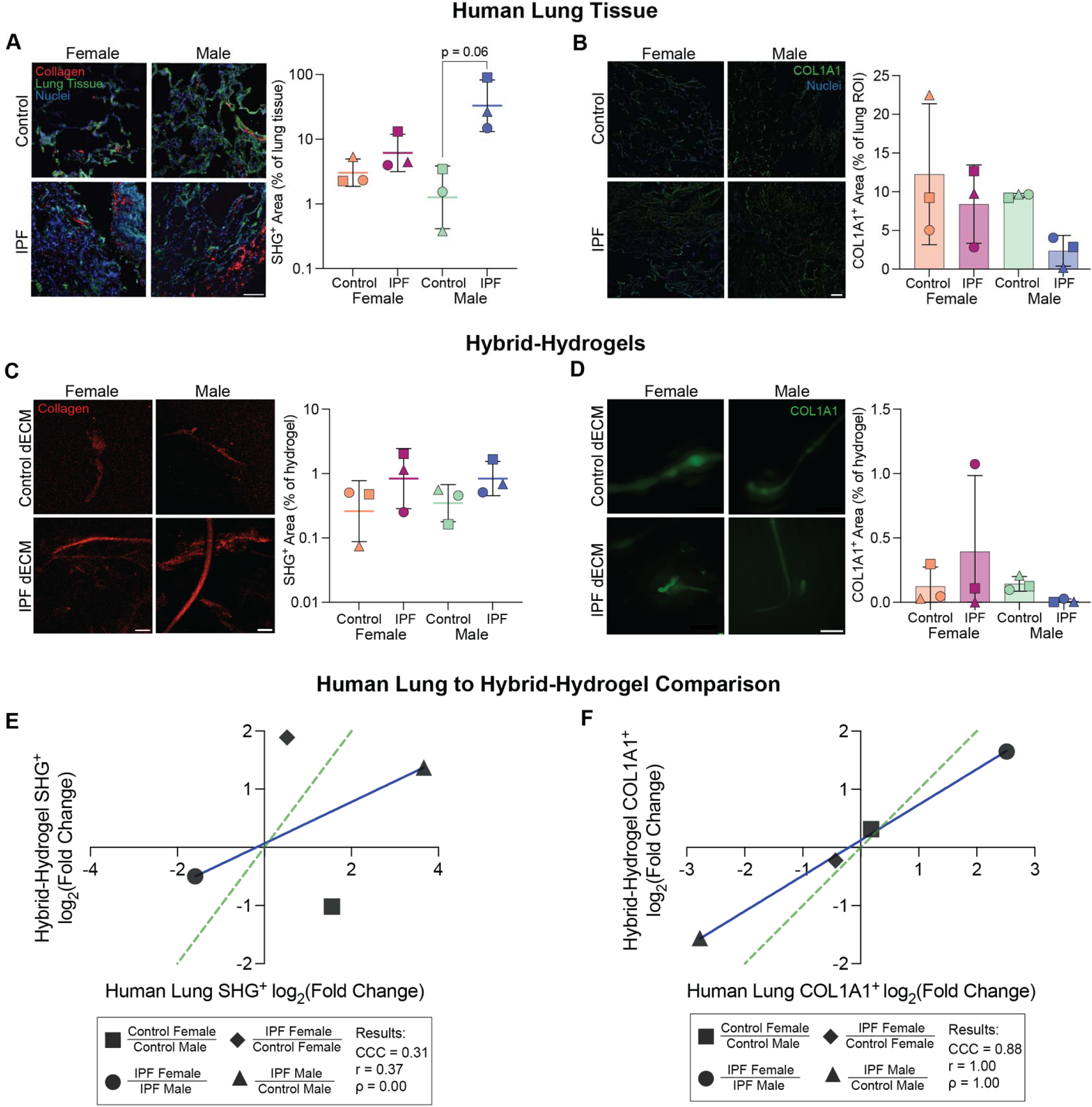
Hybrid-hydrogels retain fibrillar collagen with greater preservation of COL1A1-associated abundance patterns than higher-order collagen organization. (A) Representative SHG images and quantification of SHG-positive area in female and male control and IPF human lung tissue. (B) Representative COL1A1 immunofluorescence images and quantification of COL1A1 positive area in human lung tissue. (C) Corresponding SHG images and quantification in donor-derived hybrid-hydrogels. (D) COL1A1 immunofluorescence images and quantification in hybrid-hydrogels. (E) Human lung-to-hybrid-hydrogel comparison of SHG effects across four predefined sex- and disease-associated contrasts. (F) Corresponding comparison of COL1A1 effects. Effects are expressed as log□ fold changes; dashed lines indicate the line of identity, and solid lines represent fitted relationships. COL1A1-associated effects exhibited stronger concordance (CCC = 0.88) than SHG-defined fibrillar collagen organization (CCC = 0.31), indicating greater preservation of relative COL1A1 abundance patterns following hybrid-hydrogel fabrication.

### 3.6. Matricellular proteins are retained in hybrid-hydrogels with sex- and disease-associated differences in fidelity

We next evaluated the matricellular proteins TNC and POSTN to determine whether sex- and disease-associated biochemical patterns observed in human lung tissue were preserved following dECM processing and hybrid-hydrogel fabrication (Fig. 6). We additionally examined TGF-β1 abundance in human lung tissue and processed dECM to assess retention of a complementary matrix-associated signaling cue. Unlike fibrillar collagens, matricellular proteins primarily regulate cell adhesion, growth-factor signaling, matrix assembly, and ECM remodeling without necessarily contributing substantially to bulk matrix mass. In human lung tissue, TNC-positive area tended to decrease from control to IPF conditions in both females and males, although substantial inter-donor variability was observed (Fig. 6A). The IPF tissue analysis incorporated alveolar and fibroblastic-focus regions to provide a broader representation of the diseased distal-lung matrix used for dECM preparation. This pattern differs from previous reports describing increased TNC expression in fibrotic lung tissue, particularly within fibroblastic foci and regions of active remodeling.^65, 76^ However, because TNC expression is spatially and temporally regulated, the observed differences may reflect variation in anatomical sampling, lesion distribution, and stage of remodeling rather than a uniform reduction throughout IPF lung tissue. Experimental studies have also implicated persistent TNC in TGF-β-associated signaling and fibrosis progression, supporting its biological relevance beyond a marker of ECM remodeling.^76, 77^

**Figure 6.**
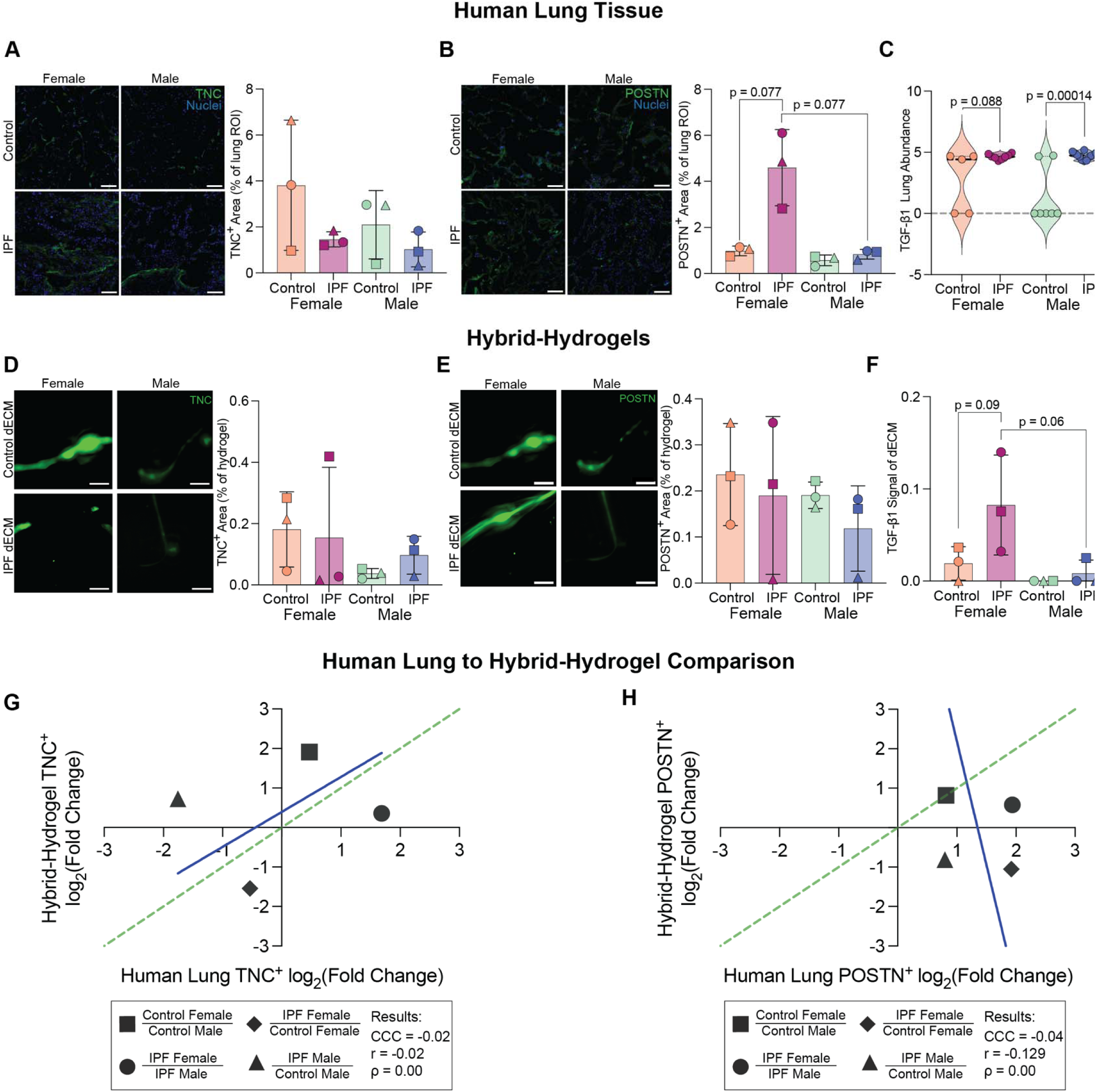
Hybrid-hydrogels retain matricellular proteins with sex- and disease-associated differences in biochemical fidelity. (A) Representative TNC immunofluorescence images and quantification of TNC-positive area in female and male control and IPF human lung tissue.. (B) Representative POSTN immunofluorescence images and quantification of POSTN-positive area in human lung tissue, showing trends toward greater POSTN-positive area in female IPF compared with female control tissue (p = 0.077) and male IPF tissue (p = 0.077). (C) Human lung proteomic abundance of TGF-β1, showing a trend toward increased abundance in female IPF relative to control (p = 0.088) and a significant increase in male IPF relative to male control (p = 0.00014). (D) Representative TNC immunofluorescence images and quantification of TNC-positive area in donor-derived hybrid-hydrogels. (E) Corresponding POSTN immunofluorescence images and quantification of POSTN-positive area in hybrid-hydrogels. (F) Relative TGF-β1 signal retained in processed human lung dECM, using a human growth factor antibody array. Female IPF-derived dECM exhibited trends toward greater TGF-β1 signal relative to female control-derived dECM (p = 0.09) and male IPF-derived dECM (p = 0.06). (G) Human lung-to-hybrid-hydrogel comparison of TNC-positive area effects (Lin’s CCC = −0.02; Pearson’s r = −0.02; Spearman’s ρ = 0.00) and (H) Corresponding comparison of POSTN-positive area effects (Lin’s CCC = −0.04; Pearson’s r = −0.129; Spearman’s ρ = 0.00) across four predefined sex- and disease-associated contrasts. Effects are expressed as log□ fold changes; dashed lines indicate the line of identity, and solid lines represent fitted relationships. Concordance statistics are interpreted descriptively because the four contrasts are group-level comparisons rather than independent paired biological samples. TNC and POSTN remained detectable in hybrid-hydrogels, with protein- and condition-specific differences in preservation of human lung abundance patterns, while TGF-β1 remained detectable in processed dECM.

Following dECM processing and hydrogel formation, TNC remained readily detectable across engineered conditions, demonstrating successful retention of this matricellular protein (Fig. 6D). The female disease-associated pattern was also retained, with a reduction from control to IPF remaining apparent in female-derived hybrid-hydrogels. In contrast, male-derived hybrid-hydrogels did not reproduce the corresponding control-to-IPF relationship observed in human lung tissue. The corresponding human lung-to-hybrid-hydrogel comparison revealed limited overall concordance across the four predefined biological contrasts (Lin’s CCC = −0.02; Pearson’s r = −0.02; Spearman’s ρ = 0.00; Fig. 6G). These results demonstrate successful TNC retention, including preservation of the female disease-associated pattern, despite incomplete agreement across all sex- and disease-associated comparisons.

POSTN showed a markedly different pattern of remodeling. Female IPF lung tissue exhibited a trend toward increased POSTN-positive area relative to female control tissue (p = 0.077) and male IPF tissue (p = 0.077; Fig. 6B), whereas the corresponding male disease-associated change was comparatively small. These observations identify a trend toward female-enriched POSTN-associated matrix remodeling, although the comparisons did not meet the conventional threshold for statistical significance. The biological relevance of this difference extends beyond POSTN as a marker of ECM accumulation. POSTN is a matricellular protein that is increased in human IPF lung, where it has been localized prominently to fibroblastic foci as well as subepithelial and subendothelial regions, and circulating POSTN has been associated with subsequent clinical progression in patients with IPF.^66^ Mechanistically, POSTN can regulate profibrotic cell–matrix signaling through integrin-associated pathways and reciprocal interactions with TGF-β signaling, while experimental disruption of POSTN-associated signaling has been shown to attenuate fibroblast proliferation and pulmonary fibrosis.^66, 78, 79^ These established functions position POSTN as a biologically relevant component of sex-associated ECM remodeling, although the present study does not establish a causal role in the observed mechanical differences.

The greater POSTN-positive area observed in female IPF lung is particularly notable because female tissue exhibited comparatively modest disease-associated mechanical stiffening (Fig. 4B). This pattern illustrates how remodeling of a biologically active matricellular cue without a corresponding increase in tissue stiffness. Considered alongside with the AFM and SHG findings, these data further distinguish the mechanical, structural, and biochemical features of sex-associated IPF remodeling: the male IPF condition was characterized by particularly pronounced mechanical and SHG-defined fibrillar collagen remodeling, whereas female IPF tissue exhibited a trend toward greater POSTN-associated biochemical remodeling. These differences do not indicate uniformly greater fibrosis in either sex but instead highlight distinct contributions of mechanical, structural, and signaling-associated remodeling to the female and male IPF microenvironments.

POSTN was also retained following dECM processing and hybrid-hydrogel fabrication, although the condition-dependent abundance patterns differed from human lung tissue. POSTN remained detectable within the engineered matrices (Fig. 6E), demonstrating retention of this additional biologically active matricellular protein. However, the female IPF-associated increase observed in human lung tissue was not reproduced, and IPF-derived hybrid-hydrogels exhibited modestly lower POSTN-positive area relative to the corresponding controls. The human lung-to-hybrid-hydrogel POSTN effect comparison further illustrated this divergence, with several engineered-material effects differing substantially in magnitude or direction from the corresponding human lung effects (Lin’s CCC = −0.04; Pearson’s r = −0.129; Spearman’s ρ = 0.00; Fig. 6H). These results demonstrate that hybrid-hydrogels retain multiple matricellular components, while preservation of condition-dependent abundance relationships varies by protein. Given the established involvement of POSTN in integrin signaling, fibroblast proliferation, and profibrotic remodeling, the altered POSTN-associated patterns identify a biochemical feature that may benefit from targeted supplementation or independent engineering into sex-informed biomaterials.^66, 78, 79^

TGF-β1 provided a complementary assessment of matrix-associated signaling beyond the matricellular proteins TNC and POSTN. Human lung proteomic analysis identified disease-associated differences in TGF-β1 abundance, including a trend toward increased abundance in female IPF relative to female control tissue (p = 0.088) and a significant difference between male IPF and male control tissue (p = 0.00014; Fig. 6C). Following decellularization and ECM processing, TGF-β1 remained detectable in donor-derived dECM using a human growth factor antibody array (Fig. 6F). Female IPF-derived dECM exhibited trends toward greater relative TGF-β1 signal compared with female control-derived dECM (p = 0.09) and male IPF-derived dECM (p = 0.06). These measurements establish retention of TGF-β1 immunoreactivity following processing, although differences in analytical methods prevent direct quantitative assessment of human lung-to hybrid-hydrogel fidelity.

Retention of TGF-β1 is biologically meaningful because the ECM acts as a reservoir for latent TGF-β complexes, which are anchored to matrix components through interactions involving latent TGF-β-binding proteins. Activation of matrix-associated TGF-β1 can depend on integrin-mediated mechanical forces and the physical properties of the surrounding ECM.^16^ However, the present growth factor antibody array confirms TGF-β1 immunoreactivity without distinguishing active from latent ligand or demonstrating retained signaling activity. Thus, retention of TGF-β1 identifies a potentially important signaling-associated component of processed dECM, although the bioavailability an activity of the retained ligand remain to be established.

The retention of TNC, POSTN, and TGF-β1 demonstrates that human lung-derived ECM contributes multiple biologically relevant signaling-associated components to sex-informed engineered biomaterials. TNC and POSTN remained detectable within hybrid-hydrogels with protein- and condition-specific differences in preservation of human lung abundance patterns, while TGF-β1 remained detectable in processed dECM. These highlight the importance of evaluating both molecular retention and condition-dependent biochemical fidelity when developing sex- and disease-informed hybrid-hydrogels.

### 3.7. Proteoglycan and GAG-rich matrix features are retained in hybrid-hydrogels with assay-dependent fidelity

Proteomic analysis also revealed pronounced remodeling of structural proteoglycans, particularly within fibroblastic foci (Fig. 3), motivating direct characterization of the GAG-rich ECM compartment. Structural proteoglycans including HSPG2, DCN, BGN, VCAN, LUM, and PRELP contribute to collagen fibrillogenesis, tissue hydration, growth-factor sequestration, and matrix viscoelasticity, making this matrix class a potential contributor to both biochemical and mechanical remodeling. Unlike fibrillar collagen, proteoglycan function depends on both core-protein abundance and GAG-chain properties, including sulfation state, charge density, hydration, and spatial organization.^36, 71^ Human IPF lung has previously been shown to exhibit increased GAG deposition and altered heparan-sulfate structure, while region- and disease-specific GAG composition persists within decellularized human lung ECM.^48, 71^

Summed structural-proteoglycan abundance from the proteomic dataset revealed sex-associated patterns of remodeling in human lung tissue. Male IPF tissue exhibited greater structural-proteoglycan abundance relative to male control tissue (p = 0.022; Fig. 7A). This disease-associated increase identifies proteoglycans as an additional component of male IPF matrix remodeling alongside the pronounced mechanical and fibrillar collagen changes described above. Because proteomics measures core-protein abundance rather than attached GAG chains, we next examined GAG-rich matrix using complementary histological and biochemical approaches. Alcian Blue staining revealed greater mean GAG-rich matrix staining in IPF tissue relative to control tissue in both sexes, with the greatest mean Alcian blue-positive area observed in female IPF samples despite substantial inter-donor variability (Fig. 7B).. In male IPF lung, increased GAG-rich matrix staining accompanied disease-associated stiffening and increased SHG-positive fibrillar collagen observed in Figures 4 and 5, highlighting remodeling across multiple ECM dimensions. Although the present measurements do not establish a causal relationship, GAG-associated changes may influence the biochemical and physical environment in which fibrillar collagen remodeling occurs.

**Figure 7.**
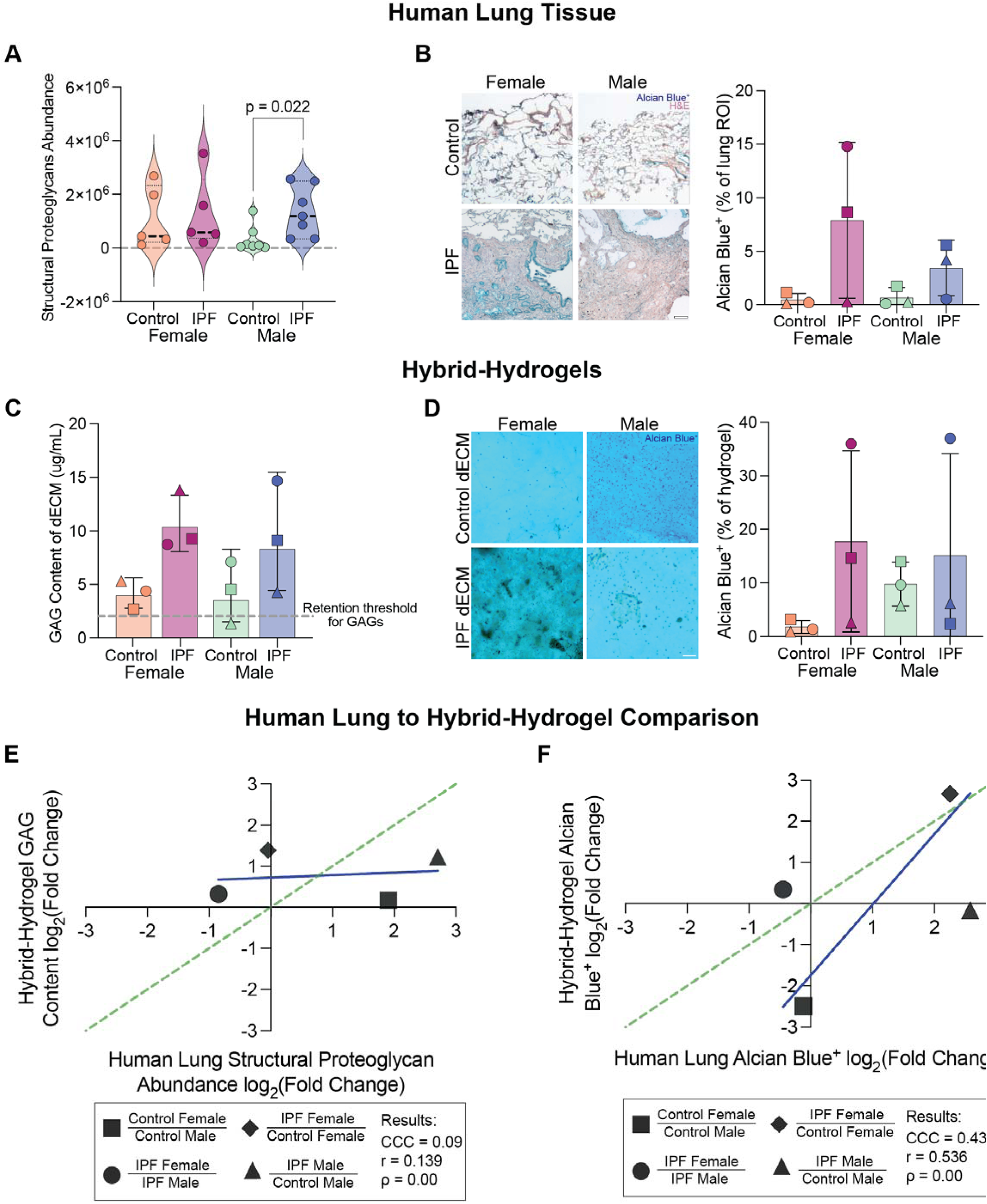
GAG-rich matrix is retained following hybrid-hydrogel fabrication with assay-dependent tissue-to-hybrid-hydrogel fidelity. A) Summed structural-proteoglycan abundance in female and male control and IPF human lung tissue measured by matrisome proteomics. Male IPF tissue exhibited greater structural-proteoglycan abundance relative to male control tissue (p = 0.022). (B) Representative Alcian Blue staining of female and male control and IPF human lung tissue and quantification of normalized Alcian Blue-positive area. IPF tissue exhibited greater mean Alcian Blue-positive area than control tissue in both sexes, with substantial inter-donor variability. (C) Total GAG concentration in processed female and male control and IPF human lung dECM, expressed as µg/mL. The dashed horizontal line indicates the GAG-retention reference threshold.^51, 80, 81^ Female IPF-derived dECM exhibited the greatest mean recovered GAG content. (D) Representative Alcian Blue staining of female and male control- and IPF-derived hybrid-hydrogels and quantification of Alcian Blue-positive area normalized to total hydrogel area. Alcian Blue-positive area was greater on average in IPF-derived hybrid-hydrogels than in the corresponding controls. (E) Cross-assay comparison of human lung structural-proteoglycan abundance and processed-dECM total GAG concentration across four predefined sex- and disease-associated contrasts (Lin’s CCC = 0.09; Pearson’s r = 0.139; Spearman’s ρ = 0.00). (F) Corresponding human lung-to-hybrid-hydrogel comparison of Alcian Blue-positive area (Lin’s CCC = 0.43; Pearson’s r = 0.536; Spearman’s ρ = 0.00). Effects are expressed as log□ fold changes; dashed lines indicate the line of identity, and solid lines represent fitted relationships. Alcian Blue comparison demonstrated modest positive tissue-to-biomaterial concordance, supporting partial preservation of GAG-rich matrix patterns in hybrid-hydrogels. Concordance statistics are interpreted descriptively because the four contrasts represent group-level comparisons rather than independent paired biological samples.

Total GAG analysis of processed dECM provided a complementary perspective (Fig. 7C). Female IPF-derived dECM exhibited the greatest mean GAG content, while male IPF-derived dECM also exhibited greater mean GAG concentration than male control-derived dECM. These results establish that measurable GAG content persists following decellularization, with disease-associated differences evident in the processed material, consistent with previous reports of region- and disease-associated variation in human lung dECM composition.^48^ However, proteoglycan core-protein abundance, total GAG concentration, and Alcian Blue-positive area interrogate distinct molecular and histochemical features of the GAG-rich matrix. Consistent with this distinction, the structural-proteoglycan-versus-total-GAG comparison exhibited negligible descriptive concordance across the four predefined biological contrasts (Lin’s CCC = 0.09; Pearson’s r = 0.139; Spearman’s ρ = 0.00; Fig. 7E). This limited agreement indicates that differences in proteoglycan core-protein abundance do not directly predict recovered GAG concentration, rather than establishing loss of proteoglycans during processing.

Following incorporation into PEG MA hybrid-hydrogels, Alcian Blue-positive matrix remained readily detectable, with greater mean positive area in IPF-derived materials than corresponding controls in both sexes (Fig. 7D). The direction of these disease-associated differences was consistent with human lung tissue, although the magnitude varied among donor-derived materials. The human lung-to-hybrid-hydrogel Alcian Blue comparison exhibited modest positive descriptive concordance (Lin’s CCC = 0.43; Pearson’s r = 0.536; Spearman’s ρ = 0.00; Fig. 7F), indicating partial preservation of the sex- and disease-associated histochemical patterns. Unlike the cross-assay comparison of proteoglycan abundance and total GAG concentration, Alcian Blue provides a directly comparable endpoint in human lung tissue and hybrid-hydrogels, strengthening interpretation of tissue-to-biomaterial fidelity.

These analyses establish that GAG-rich matrix features remain detectable following dECM processing and hybrid-hydrogel fabrication, with disease-associated differences retained in the engineered biomaterials. Importantly, the extent of correspondence depends on the matrix property and analytical approach: proteomics measures core-protein abundance, biochemical assays quantify recovered GAG concentration, and Alcian Blue staining measures the fraction of tissue or hybrid-hydrogel area containing acidic polysaccharide-rich matrix. The partial preservation of human lung Alician Blue patterns identifies GAG-rich matrix as a tissue-informed design criterion that can be retained in hybrid-hydrogels while highlighting opportunities to further refine biochemical fidelity through targeted biomaterial engineering approaches.

## 4. Conclusion

This study establishes a tissue-informed framework for engineering sex- and disease-specific biomaterials by integrating region-resolved human lung matrisome proteomics with complementary mechanical, structural, histological, and biochemical characterization. Female and male IPF lungs exhibited distinct patterns of ECM remodeling across anatomical regions and disease-associated states, extending beyond differences in collagen abundance or tissue stiffness alone. Notably, male IPF lungs exhibited pronounced disease-associated tissue stiffening and fibrillar collagen remodeling, while female IPF tissue showed comparatively modest mechanical changes despite evidence of biochemical remodeling, including differences in matricellular proteins and proteoglycan/GAG-rich matrix. These findings highlight the importance of defining sex-associated ECM microenvironments through multiple complementary measurements rather than a single indicator of fibrosis.

Translation of these tissue-derived design criteria into PEGαMA hybrid-hydrogels revealed feature-specific differences in tissue-to-biomaterial fidelity. Controlled hydrogel formulations closely reproduced sex- and disease-associated mechanical relationships observed in human lung tissue, while fibrillar collagen organization, matricellular protein abundance, and GAG-rich matrix showed varying degrees of preservation following dECM processing and hybrid-hydrogel fabrication. In particular, COL1A1-associated abundance patterns were more consistently reproduced than SHG-defined fibrillar collagen features, while TNC and POSTN remained detectable despite limited preservation of condition-dependent abundance patterns. Alcian Blue-positive area demonstrated partial human lung-to-hybrid-hydrogel correspondence, while TGF-β1 remained immunoreactive in processed dECM. These findings distinguish biochemical retention from preservation of biologically relevant matrix properties and demonstrate the importance of validating individual ECM features following dECM processing and hybrid-hydrogel fabrication.

The pronounced male predominance of IPF highlights a critical need to understand how biological sex influences disease susceptibility and progression. Our findings identify distinct extracellular microenvironments in female and male human lungs, establishing ECM composition, organization, and mechanics as important dimensions of sex-associated fibrotic remodeling. By translating these differences into tissue-informed biomaterial design criteria and identifying which features are retained or require additional engineering, this study provides a foundation for investigating how sex-associated ECM cues regulate cellular responses to injury and fibrosis. This framework moves beyond conventional disease-matched biomaterials toward sex- and disease-informed human models capable of uncovering mechanisms underlying divergent disease trajectories, informing targeted therapeutic strategies for IPF, and advancing the study of other conditions characterized by sex-associated differences in susceptibility and progression.

## Acknowledgements

Human lung samples were provided by the Human Tissue and Cell Core at the University of Colorado Anschutz, supported by NIH-NHLBI grant P01HL162607. Mass Spectrometry was performed by the Mass Spectrometry Proteomics Shared Resource Facility [RRID SCR_021988] at the University of Colorado Anschutz. Additional funding was provided by the National Heart, Lung, and Blood Institute HL153096 (HN, RB, MCM, DE, CMM), National Science Foundation 1941401 (MCM, CMM), and NIH/NCATS Colorado CTSA Grant Number T32 TR004367, UM1 TR004399 (HN).

